# *ROSIE*: AI generation of multiplex immunofluorescence staining from histopathology images

**DOI:** 10.1101/2024.11.10.622859

**Authors:** Eric Wu, Matthew Bieniosek, Zhenqin Wu, Nitya Thakkar, Gregory W. Charville, Ahmad Makky, Christian Schürch, Jeroen R Huyghe, Ulrike Peters, Christopher I Li, Li Li, Hannah Giba, Vivek Behera, Arjun Raman, Alexandro E. Trevino, Aaron T. Mayer, James Zou

## Abstract

Hematoxylin and eosin (H&E) is a common and inexpensive histopathology assay. Though widely used and information-rich, it cannot directly inform about specific molecular markers, which require additional experiments to assess. To address this gap, we present *ROSIE,* a deep-learning framework that computationally imputes the expression and localization of dozens of proteins from H&E images. Our model is trained on a dataset of over 1000 paired and aligned H&E and multiplex immunofluorescence (mIF) samples from 20 tissues and disease conditions, spanning over 16 million cells. Validation of our *in silico mIF* staining method on held-out H&E samples demonstrates that the predicted biomarkers are effective in identifying cell phenotypes, particularly distinguishing lymphocytes such as B cells and T cells, which are not readily discernible with H&E staining alone. Additionally, *ROSIE* facilitates the robust identification of stromal and epithelial microenvironments and immune cell subtypes like tumor-infiltrating lymphocytes (TILs), which are important for understanding tumor-immune interactions and can help inform treatment strategies in cancer research.

## Introduction

H&E staining is ubiquitously used in clinical histopathology due to its affordability, accessibility, and effectiveness for discerning clinically relevant features. While H&E readily identifies nuclear and cytoplasmic morphology, its utility is limited in revealing more complex molecular information associated with modern precision medicine (*1*). Pathologists can identify diverse cell types from H&E staining alone; however, computational approaches to annotating H&E images only distinguish a few broad cell categories, such as endothelial, epithelial, stromal, and immune cells (*2–4*). These methods are valuable for detecting tumors and identifying basic structural features but are limited in revealing detailed aspects of the cellular microenvironment, such as protein expression profiles, disease signatures, or the specific identity of immune cells like lymphocytes.

In contrast, multi-plex immunofluorescence (mIF) imaging techniques such as Co-Detection by Indexing (CODEX) and immunohistochemistry (IHC) enable in situ detection of dozens of proteins simultaneously. This capability allows for the exploration of richer tissue microenvironments, offering insights that are unattainable through H&E staining alone (*5–8*). However, the application of CODEX and similar mIF techniques is limited by high costs, time-intensive protocols, and lack of adoption in clinical labs, making them less feasible for routine use (*9*).

In this work, we present *ROSIE* (**RO**bust in **S**ilico **I**mmunofluorescence from H&**E** images), a framework for *in silico* mIF staining based on an H&E-stained input image. We train a deep learning model on a dataset of over 1,000 tissue samples co-stained with H&E and CODEX. This dataset, comprising nearly 30 million cells, is the largest of its kind to date and significantly surpasses the scale of previous studies, which typically focus on data from a single clinical site or a limited number of stains. Our findings demonstrate that the proposed method can robustly predict and spatially resolve dozens of proteins from H&E stains alone.

We validate the biological accuracy of these *in silico-generated* protein expressions by employing them in detailed cell phenotyping and the discovery of tissue structures such as stromal and epithelial tissues. Our approach enables the identification of immune cell subtypes, including B cells and T cell subtypes that are not discernible by H&E staining alone, thus offering a powerful tool for enhancing the diagnostic and research potential of standard histopathological practices.

### Related works

Recent advancements in training histopathology foundation models (*10–12*) have demonstrated that models trained on large, diverse sets of histology images in an unsupervised manner can yield strong performance when adapted to downstream tasks like predicting tissue types and disease diagnosis and prognosis. While foundation models can learn intricate biological features within the distribution of H&E images, they still need to be explicitly trained on other imaging modalities and molecular information to be adapted for generative methods like *in silico* staining.

Previous works in predicting immunostains from H&E have typically focused on small paired or unpaired datasets and imputing up to several biomarkers at once. To start, VirtualMultiplexer is a GAN-based method for predicting 6-plex IHC stains (*13*) using unpaired H&E and IHC samples. The limitation of using unpaired samples is that validation of predictions is limited to qualitative or visual assessments. Several methods have been trained on paired (adjacent slice) datasets, such as: Multi-V-Stain (*14*)), which predicts a 10-plex mIMC panel on 336 melanoma samples; DeepLIIF (*15*) which predicts 3-plex mIHC on one sample; and other GAN-based methods for predicting single or several biomarkers (*16*), (*17–21*). (*22*) predict two IHC biomarkers on brain tissue. Compared with paired samples, co-stained (or same slice) samples allow for direct pixel-level alignment and prediction from H&E to immunostain; to this end, HEMIT (3-plex mIHC) (*23*) and vIHC (1-plex mIHC) both are trained on co-stained samples but are also limited to evaluation on a single sample. (*24*) focus on predicting a transcriptomics panel (1000 genes) using 4 co-stained samples. Specific to multiplexed immunofluorescence, 7-UP (*25*) used a small 7-plex panel to predict over 30 biomarkers. (*26*) use autofluorescence and DAPI channels to infer seven biomarkers.

Our method improves upon this previous body of work in several ways. First, we train and evaluate the largest co-stained H&E and immunostaining dataset with over 1000 samples. Second, whereas previous datasets were limited to one or several tissue types, our dataset spans ten body areas and disease types. Third, whereas previous works focus on visual or quantitative metrics of expression prediction only, ours demonstrates the usefulness of the predicted expressions for cell phenotyping and tissue structure discovery. Finally, instead of using difficult-to-train adversarial methods, we use a straightforward single MSE objective for training our model.

## Results

### A comprehensive, diverse dataset of co-stained tissue samples

We introduce a first-of-its-kind training and evaluation dataset of 23 studies spanning nearly 30M cells, over 2000 samples, and 259M unique patches (Figure 1A, Table 1; see Supplementary Materials for dataset details). All studies are on tissue samples with H&E and CODEX co-staining on the exact same samples. We hold out four studies for evaluation while training and validating with the remaining nineteen. All datasets consist of tissue microarray (TMA) cores (average 10K cells per core) except UChicago-DLBCL, which contains full slide samples (average 1.5M cells per slide). Stanford-PGC is a study of patients with pancreatic and gastrointestinal cancer from Stanford Healthcare. Ochsner-CRC is a study of patients with colorectal cancer from Ochsner Medical Center. Tuebingen-GEJ is a study of patients with cancer in the gastroesophageal junction from Tübingen University Hospital. UChicago-DLBCL is a study of patients with diffuse large B-cell lymphoma from the University of Chicago Medical Center.

**Fig. 1.**
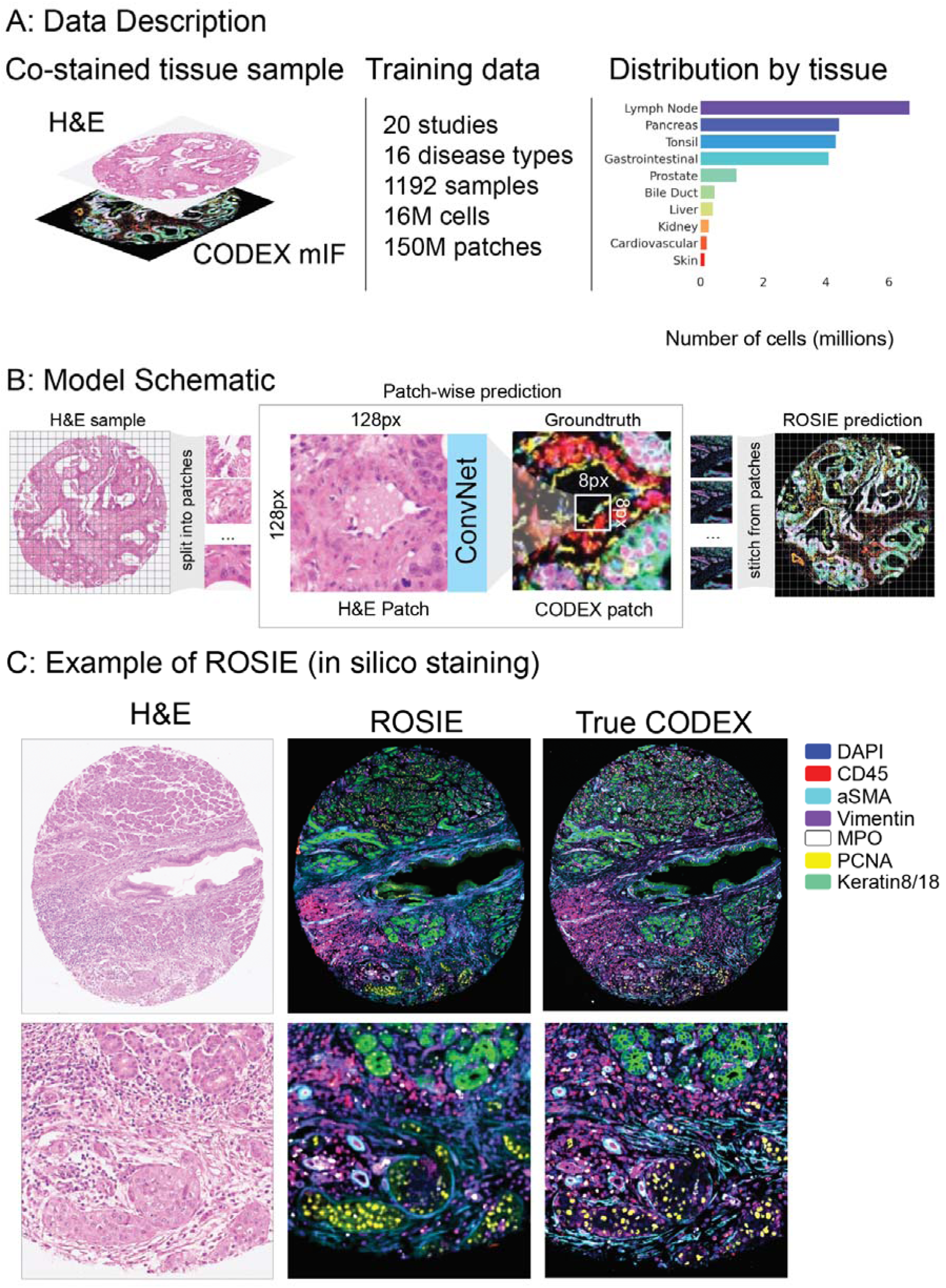
Overview of *ROSIE*. **(A)** Our training dataset consists of 20 studies and over 16M cells. Each tissue sample is co-stained with H&E and CODEX. 16 disease types and 10 body areas are represented in this dataset. The overall distribution of represented tissue types across training and evaluation datasets is shown on the right. (**B**) Given an H&E-stained image, *ROSIE* predicts the pixel-level expression of 50 biomarkers. An exemplar image is visualized, where seven representative biomarkers are colored and shown alongside the true CODEX image. While the generated images used in our analyses are produced with 8px striding, this image is produced using 1px striding for greater visual clarity. (**C)** A schematic of model training and inference is shown. Given an H&E sample, the image is split into patches of size 128px by 128px. The model is trained to predict the average expressions of the center 8px by 8px patch in the corresponding CODEX image. After the model is trained, a predicted CODEX image is generated by aggregating all of the generated patches into a single image.

**Table 1.** Description of evaluation datasets. . We describe the four evaluation datasets used in this study. Samples from Stanford-PGC and UChicago-DLBCL are divided into training and test splits; on the other hand, no samples and disease types from Ochsner-CRC and Tuebingen-GEJ are used in the model’s training data. UChicago-DLBCL contains two full tissue samples, whereas the rest of the datasets consist of TMA core samples.

| Evaluation Datasets | In training data? |  | # Samples | # Slides | # Cells |
| --- | --- | --- | --- | --- | --- |
|  | Study | Disease |  |  |  |
| Stanford-PGC | Yes | Yes | 149 | 1 | 817,765 |
| Ochsner-CRC | No | No | 94 | 1 | 635,649 |
| Tuebingen-GEJ | No | No | 240 | 1 | 365,734 |
| UChicago-DLBCL | Yes | Yes | 2 | 1 | 3,099,419 |
| Total | - | - | 485 | 8 | 4,918,567 |

### Generative deep learning model for inferring protein expression from H&E stains

*ROSIE* is a framework for *in silico* staining on a sample based on an H&E image. Using a ConvNext (*27*) convolutional neural network (CNN) architecture, *ROSIE* operates on the patch level: given an input 128x128 pixel patch, it produces a prediction for the average expressions of the biomarker panel across the center 8x8 pixels (Figure 1B). Using a sliding window with an 8px step size, we iteratively generate predictions on all 8x8 pixel patches within a sample, then stitch together predictions to produce a whole, contiguous image. *ROSIE* can be run with smaller sliding window sizes, down to 1px for native resolution outputs (see Figure S6 for examples). Due to computational tradeoffs, analyses performed in this paper are on the standard 8px sliding window setting. Although Vision Transformer (ViT) models (*28*) have recently gained traction as top-performing histopathology foundation models, we find that ConvNext outperforms several ViT models despite its smaller size (Table S1).

A total of 148 unique biomarkers are represented across all studies. We constrain our method to predict the top 50 biomarkers by prevalence. While all evaluation studies are stained with these 50 biomarkers, some studies used in training do not; in these cases, only the subset of biomarkers present in this set are used. The full biomarker set that the model is trained to predict includes (in order of prevalence):

*DAPI, CD45, CD68, CD14, PD1, FoxP3, CD8, HLA-DR, PanCK, CD3e, CD4, aSMA, CD31, Vimentin, CD45RO, Ki67, CD20, CD11c, Podoplanin, PDL1, GranzymeB, CD38, CD141, CD21, CD163, BCL2, LAG3, EpCAM, CD44, ICOS, GATA3, Gal3, CD39, CD34, TIGIT, ECad, CD40, VISTA, HLA-A, MPO, PCNA, ATM, TP63, IFNg, Keratin8/18, IDO1, CD79a, HLA-E, CollagenIV, CD66*

### *ROSIE* accurately predicts protein biomarker expressions

When applying *ROSIE* to the four evaluation datasets, we report a Pearson R correlation of 0.285, a Spearman R correlation of 0.352, and a sample-level C-index of 0.706 when comparing the ground truth and computationally generated expressions across all 50 biomarkers in all four datasets (Table 2). Whereas the Pearson correlation indicates a linear predictive relationship, the Spearman R and C-index indicate the usefulness of the predicted expressions in clinical tasks that involve ordering cells or samples by expressing a certain biomarker (e.g., identifying immune markers in a cancer patient cohort). C-index refers to the concordance index computed on the sample level using the 75th percentile expression value as a threshold. A C-index of 0.5, for instance, indicates random chance. We show that our method significantly outperforms two baseline methods: *H&E expression*, which uses the average intensity across RGB channels as a direct proxy for protein expression for every biomarker, is intended to test whether our predictive accuracy is simply due to recapitulating the staining signal; and *cell morphology,* which uses morphology features derived from the cell segmentations in addition to the three RGB channels as inputs to a multi-layer perceptron (MLP) neural network trained to predict protein expression, is intended as a representative machine learning model that uses common H&E-derived features as input. Both baseline methods reported near-at-random performance based on the three evaluation metrics. Figure 1C visualizes an exemplar predicted sample with a representative seven biomarker panel.

**Table 2.** Main evaluation results. . Our method is evaluated using three metrics: Pearson R, Spearman R, and C-index (sample), which refers to the concordance index computed on the sample level using the 75th percentile of expression values in a sample as a threshold. The performance of our method on the primary dataset (Stanford-PGC) is reported along with two baseline methods: H&E expression, which uses the mean cell-wise H&E pixel value as a proxy for protein expression, and cell morphology, which uses features derived from the cell outlines based on DAPI expression as well as the RGB pixel values as input to a neural network to predict protein expression. We also report the performance of our method on the other three evaluation datasets.

| Evaluation Datasets (50 biomarkers) | Pearson R | Spearman R | C-index (sample) |
| --- | --- | --- | --- |
| Stanford-PGC |  |  |  |
| - H&E expression | 0.007 | 0.013 | 0.504 |
| - Cell morphology | 0.072 | 0.081 | 0.574 |
| - **ROSIE (our method)** | 0.319 | 0.386 | 0.694 |
| ROSIE evaluated on: |  |  |  |
| - Ochsner-CRC | 0.218 | 0.276 | 0.597 |
| - Tuebingen-GEJ | 0.265 | 0.289 | 0.668 |
| - UChicago-DLBCL | 0.254 | 0.327 | 0.820 |
| *Average (ROSIE)* | *0.246* | *0.297* | *0.695* |

*ROSIE* generates highly accurate full-sample CODEX images and recapitulates salient visual features across representative immune and structural biomarker panels (Figure 2A). To illustrate the robustness of *ROSIE* across a range of predictions including relatively low-performing ones, we display side-by-side predicted and ground truth samples drawn from the 99th, 75th, 50th, and 25th percentiles of performance (by Pearson R) in the Stanford-PGC dataset. Figure 2B shows the Pearson R score for each of the 50 predicted biomarkers averaged across all evaluation datasets. We also visualize every single biomarker individually (Figure S1) from Stanford-PGC (median samples by Pearson R), along with their distributions by Pearson R (Figure S2) and scores on rank tests (Spearman R and C-index, Figure S3).

**Fig 2.**
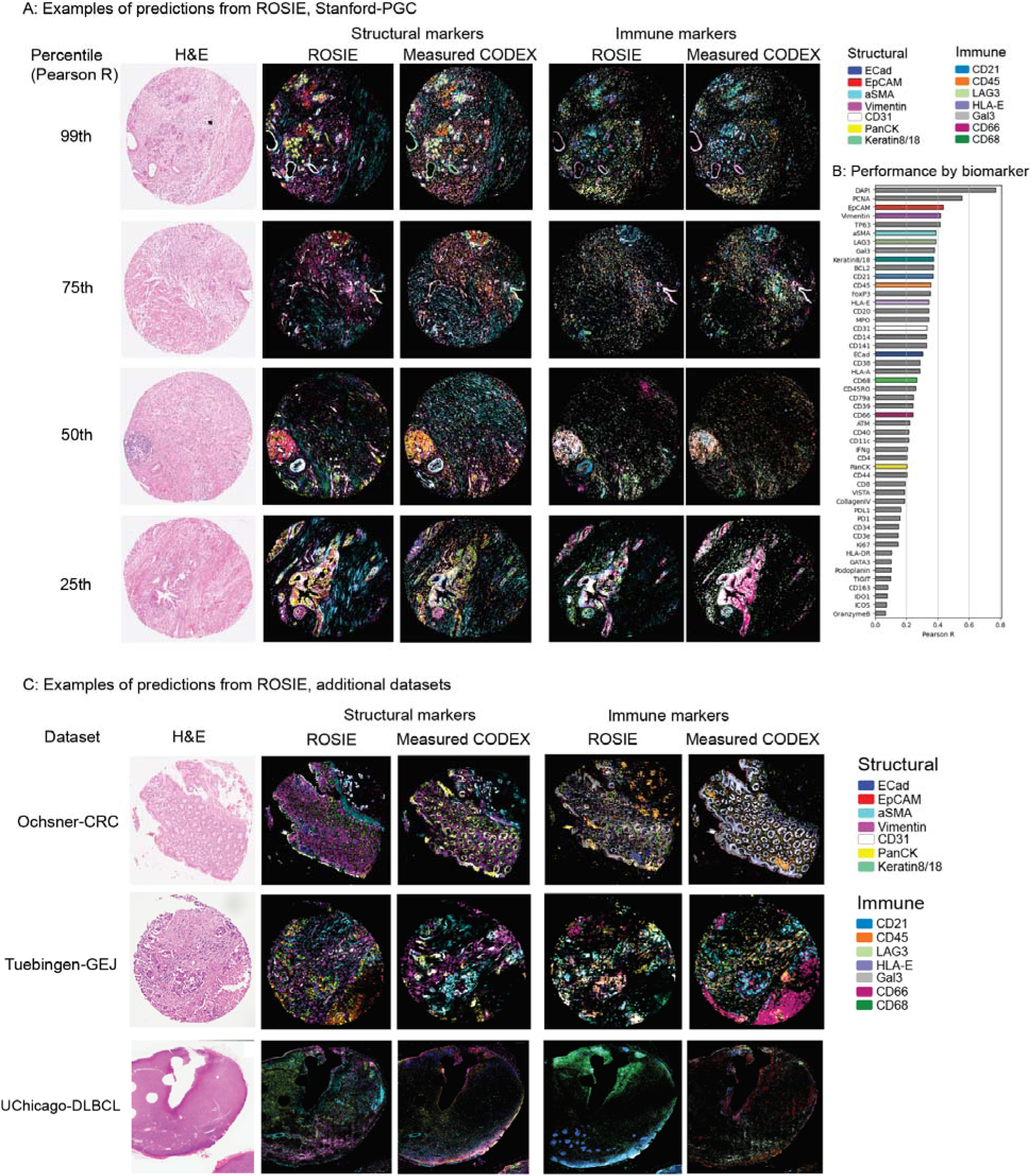
Visualization of predictions from *ROSIE*. **(A)** Predicted and measured CODEX samples along with the co-stained H&E images. The 99th, 75th, 50th, and 25th percentile (by Pearson R) samples are shown, colored with seven structural (Keratin8/18, EpCAM, Vimentin, ECad, aSMA, CD31, PanCK) and immune (HLA-E, Gal3, CD45, CD21, LAG3, CD66, CD68) markers. **(B)** Pearson R correlation on all evaluation datasets for 50 biomarkers (visualized biomarkers are colored). Figure S5 visualizes zoomed-in patches of predictions. **(C)** Visualizations of median samples (by Pearson R) from three additional datasets: Ochsner-CRC, Tuebingen-GEJ, and UChicago-DLBCL.

### Generalization to unseen studies and disease types

*ROSIE* performs robustly even when evaluated on samples from clinical sites and disease types never seen during training. When evaluated on Ochsner-CRC and Tubingen-GEJ, two studies whose samples and disease types do not appear in the training dataset, *ROSIE* reports comparable average performance to Stanford-PGC: a Pearson R of 0.241 (vs. 0.319 on Stanford-PGC), Spearman R of 0.283 (vs. 0.386 on Stanford-PGC), and a sample-level C-index of 0.633 (vs. 0.694 on Stanford-PGC) across all 50 biomarkers (see Table 2). We confirm these results visually in Figure 2C which contains median samples (by Pearson R) from three other evaluation datasets (Ochsner-CRC, Tuebingen-GEJ, and UChicago-DLBCL) with the same representative immune and structural biomarker panels.

### A simple metric for postprocessing and filtering *in silico* staining quality

Batch effects, or variations across histopathology samples due to factors like staining quality, tissue type, and artifacts, can significantly affect deep learning model generalizability (*29*, *30*). It is desirable, therefore, to be able to predict which samples might be of lower quality due to batch effects and exclude them from downstream analyses.

To this end, we introduce two simple but effective heuristics for scoring predicted samples on staining quality: dynamic range, which is a measure of the difference between the 99th and 1st percentile values in a biomarker stain; and W1 distance, which is the average Wasserstein distance between a test H&E image’s histogram distribution and all histogram distributions from the training H&E image dataset. Figure S6 shows the effect of applying each quality filter to the four evaluation datasets: using the median as a cutoff for out-of-distribution samples, the average Pearson R score increases from 0.285 to 0.312 using W1 distance and to 0.336 using the dynamic range. Panel A of Figure S6 illustrates the relationship between the dynamic range and Pearson R score. Panel B likewise shows the relationship between the predicted Wasserstein distance and Pearson R score.

### Biomarker predictions are useful for cell and tissue phenotyping

Given that the protein biomarker expressions generated by *ROSIE* are highly correlated with ground truth measurements, we validate their biological and clinical usefulness by using them in phenotyping cells. To do this, we first train a nearest-neighbor algorithm to predict annotated cell labels on the ground truth CODEX biomarker expressions. Then, we input the biomarker expressions generated by *ROSIE* into the algorithm to produce cell type predictions. *ROSIE* can predict seven cell types (B cells, Endothelial cells, Epithelial cells, Fibroblasts, Macrophages, Neutrophils, and T cells) significantly better than a model using cell morphology and H&E RGB channels as inputs (*cell morphology)* or randomly assigning cell types according to average sample proportions (*bulk phenotyping)* (Figure 3). Further analyses show strong B and T cell differentiation from the cell type classification confusion matrix (Figure S4). Furthermore, we perform cell phenotyping on the Ochsner-CRC dataset and find that the labels produced by the *ROSIE-*generated biomarkers performed comparably to Stanford-PGC (average F1 of 0.411 vs. 0.507, respectively) (Table S2 and Figure S4).

**Fig. 3.**
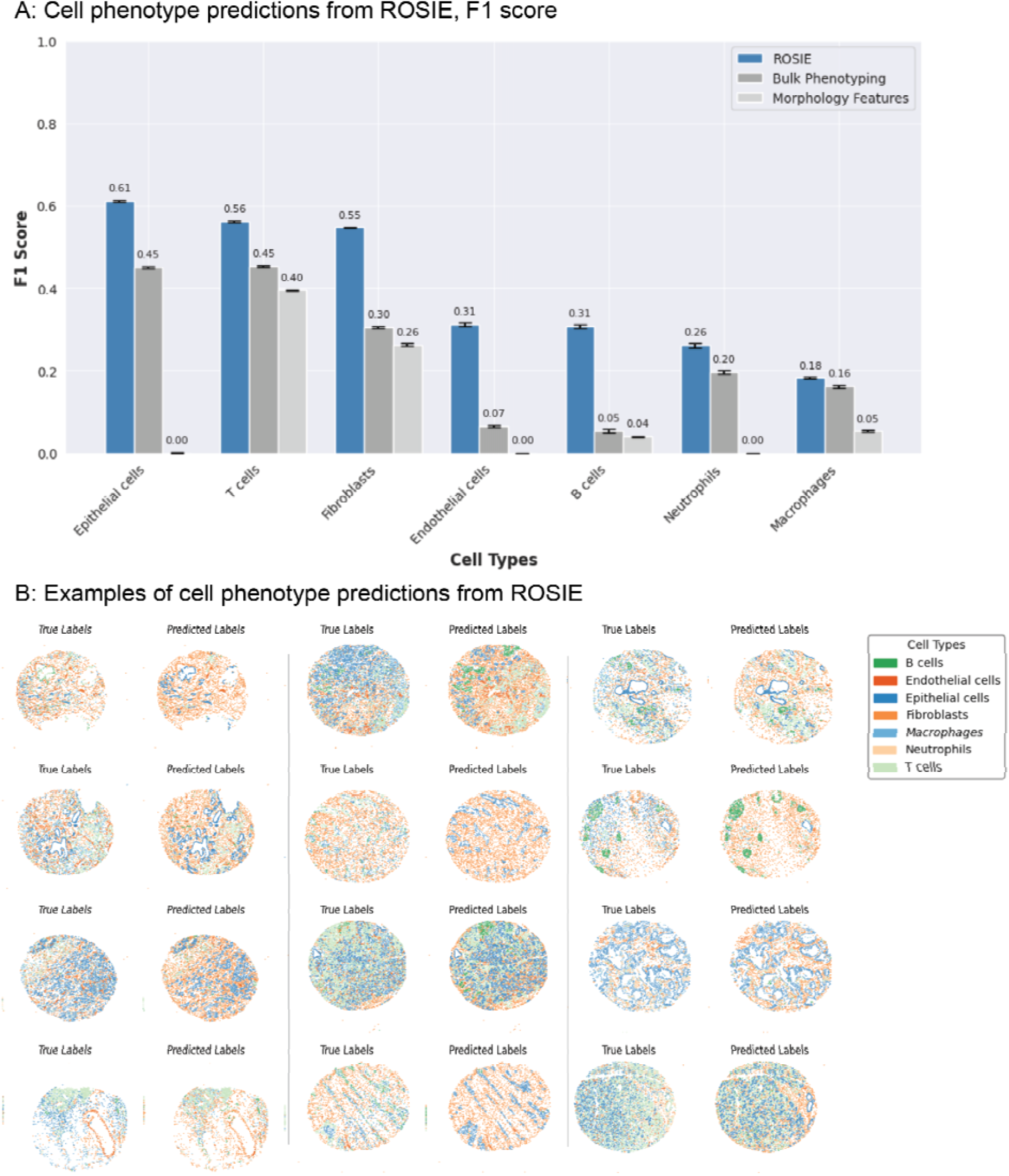
Cell type predictions using *ROSIE*. **(A)** F1 scores on the primary Stanford-PGC dataset, comparing the performance of *ROSIE* to two baselines: *bulk phenotyping*, which randomly assigns cell types based on sample-level cell type proportions, and *morphology features*, which uses a three-layer neural network to classify cells based on morphology features and the H&E RGB channels. Error bars are the 95% bootstrapped confidence intervals. **(B)** Visualization of cell phenotype predictions from twelve median samples by Pearson R.

We are interested in whether these predictions validate therapeutically relevant distinctions between tissues. Different tissue types are typically known to be immunologically “hot”, indicating greater immune cell presence and infiltration, or “cold”, implying less immune cell activity. For instance, colorectal cancer (e.g. Stanford-PGC dataset) is known to be “cold” while pancreatic cancer (e.g. Ochsner-CRC dataset) is known to be “hot” (*31*, *32*). Indeed, our results reflect this immunological validity check, with the immunologically “colder” Stanford-PGC having a lower predicted average proportion of T cells per sample of 20.0% (vs. 21.1% ground truth) and the immunologically “hotter” Ochsner-CRC having 40.1% T cells per sample (vs. 30.6% ground truth).

We also extend our method beyond single-cell phenotyping and demonstrate its effectiveness in identifying tissue structures within a sample. We use a top-performing tissue structure identification algorithm, SCGP (*33*), which projects the acquired tissue sample into a graph, with nodes as cells and edges as neighboring cell pairs. Using this graph structure, the algorithm performs unsupervised clustering to discover tissue structures based on the ground truth and *ROSIE*-generated biomarker expressions separately. Figure 4 shows structures discovered in several samples and the reported Adjusted Rand Index (ARI) and F1 scores by comparing the ground truth and generated expressions. Our method achieves average ARI and F1 scores of 0.475 and 0.624, respectively, and is significantly higher than a baseline method of expressions generated from a three-layer neural network using cell morphology features derived from cell segmentations and mean H&E RGB values as inputs (ARI of 0.105 and F1 of 0.229). Full scores are reported in Figure S8.

**Fig. 4.**
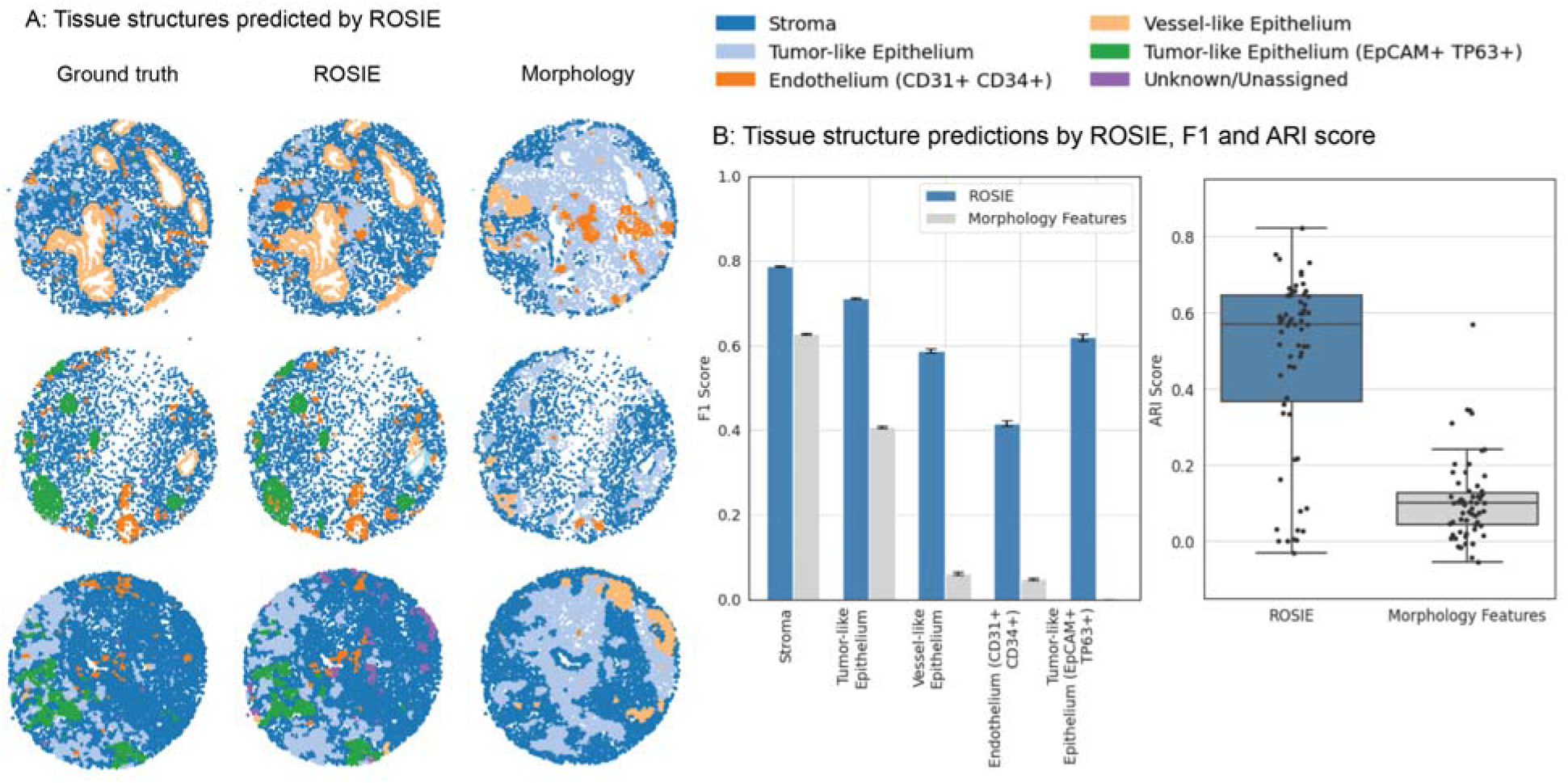
Tissue structure discovery by *ROSIE*. Discovery of tissue structures using biomarkers generated by *ROSIE* on the Stanford-PGC test dataset. Five tissue structures are identified using a graph partitioning algorithm that clusters cells based on their expression profiles and neighboring cells. This algorithm is performed on both the ground truth measured and *ROSIE-* generated biomarker expressions and then reconciled to a common label set. **(A)** visualizes several representative samples of tissue structures discovered using the ground truth CODEX measurements, *ROSIE-generated* expressions, and morphology baseline method. **(B)** We report the F1 score by comparing the structures discovered using ground truth, *ROSIE*-generated biomarkers, and morphology features. Error bars are the 95% bootstrapped confidence intervals. ARI score is also reported by comparing the unlabeled discovered clusters, where each dot is a sample.

Additionally, we use the *ROSIE-*generated expressions to identify two cell neighborhood phenotypes of interest: tumor-infiltrating lymphocytes (TILs) and lymphocyte neighboring epithelial cells (LNEs). These cell types are defined by their cellular niche and their biomarker expression profile: TILs are lymphocytes that reside in epithelial tissues, and LNEs are epithelial cells that neighbor lymphocytes. Figure 5 shows that the predicted proportions of TILs and LNEs per sample in the Stanford-GPC test dataset are highly correlated with the ground truth-derived proportions (Pearson R of 0.805 and 0.598 and Spearman R of 0.329 and 0.575 for TILs and LNEs, respectively), suggesting that the *ROSIE*-generated expressions may be useful for clinical tasks that involve estimating or ordering samples in a patient cohort by the presence of specific biomarkers, cells, or cell interactions.

**Fig. 5.**
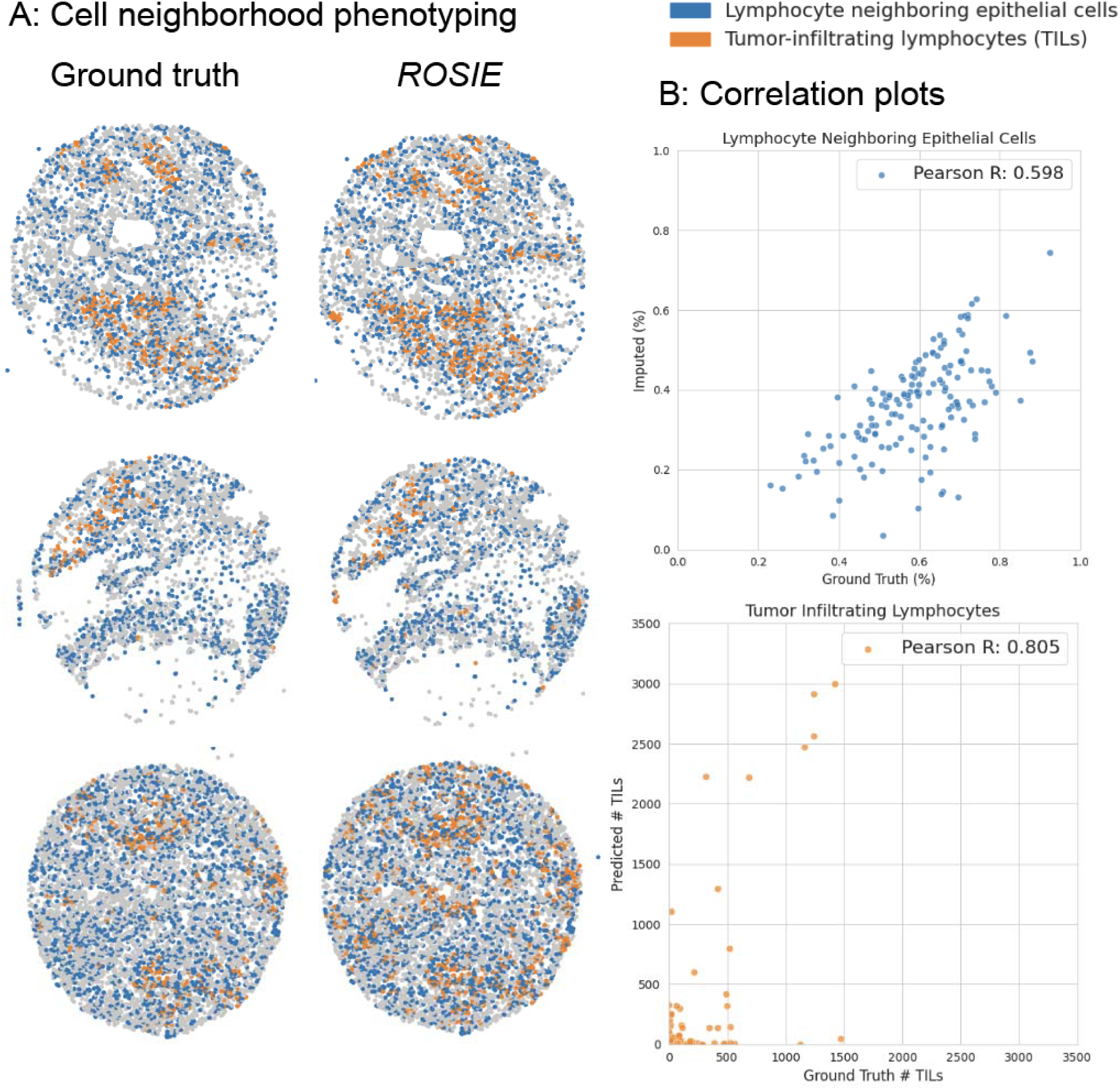
Cell neighborhood phenotyping by *ROSIE*. We identify two cellular neighborhood phenotypes of interest: tumor-infiltrating lymphocytes (TILs) and lymphocyte neighboring epithelial cells (LNEs). TILs are defined as cells that are labeled as lymphocytes (B and T cells) and reside in epithelial tissue (using the graph partitioning algorithm). LNEs are epithelial cells that have at least one lymphocyte as a neighbor. TILs are measured as the raw count per sample, whereas LNEs are measured as the proportion of epithelial cells with lymphocyte neighbors. **(A)** We visualize three samples (by median Pearson R) of TILs and LNEs based on ground truth-derived and *ROSIE*-predicted expressions. **(B)** Scatter plots of predicted and ground truth measurements, where each dot represents a sample.

## Discussion

Our study aims to bridge the gap between abundant, inexpensive H&E staining and the rich but more costly molecular information provided by multiplex immunofluorescence (mIF) staining. The primary question we shed light on is the extent to which H&E stains embed molecular hallmarks and features that could be computationally extracted. Our results indicate that although H&E staining has been traditionally limited in identifying few cell phenotypes, its structural and morphological features indeed contain significant information about protein expressions when analyzed with deep learning generative AI methods. This suggests that H&E staining has unrealized potential for use in clinical decisions that traditionally require more complex and expensive assays. Although prior work has focused on imputing up to several markers at a time, our study is the first to use deep learning to learn the relationship between H&E and up to 50 protein biomarkers at once. This setting now enables a more comprehensive view of the tissue microenvironment and offers more nuanced insights into specific tumor and immune cellular phenotypes.

Recent advances in Transformer architecture-based foundation models trained on histopathology images (*10*, *12*, *34*) show promise when adapted and fine-tuned on a wide range of clinical downstream tasks. However, our results indicate that such models still underperform our significantly smaller convolutional neural network. For instance, a ViT-L/16 Transformer model with over 300M parameters pre-trained on histopathology images still did not perform as well as ConvNext, a 50M parameter CNN pre-trained on non-pathology images. We offer several potential explanations for this counterintuitive finding: First, the inductive biases of convolutional operations in CNNs are better suited for extracting local contextual features from patch-based histology imaging; second, larger foundation models are harder to train and may overfit easier to the training data—indeed, we observed that the performance of the larger Vision Transformer models plateaus earlier in the process than compared to a CNN. This result suggests the importance of future research in analyzing when large foundation models are more appropriate than smaller CNNs.

One focus of our study is on the rank correlation of our generated expressions and cellular phenotypes. Strong rank correlations suggest that the generated biomarkers are useful in clinical settings where the relative ordering of the presence of a biomarker or phenotype is important, e.g., finding patients that are the most receptive to a therapy, or predicting patient prognosis based on a specific biomarker. For instance, we observe that while our model can effectively predict cell neighborhood phenotypes (TILs and lymphocyte neighboring epithelial cells), these predictions also exhibit biases in over-or under-predicting the proportion or counts of these phenotypes. Despite these biases, the relative ordering of these phenotypes is still largely maintained and thus is still useful in the settings mentioned above.

Inter-batch effects due to staining technique, quality, and machinery are known to cause variations in the image statistics of H&E stains. Since predictions generated on H&E stains that significantly deviate from the training data are expected to perform worse, we propose two methods for quantifying the quality of generated mIF stains. We demonstrate that computing the dynamic range of predicted expressions and calculating the Wasserstein distance between training and generated data image histograms both correlate well with the empirical prediction accuracy. We believe these can be valuable tools that accompany our deep learning framework, allowing users to determine their clinically acceptable range of stain generation quality.

We acknowledge several limitations with our data and framework. Though we perform quality control on each sample (see Supplementary Methods), the alignment of H&E and CODEX images is susceptible to artifacts such as fraying, which can lead to misalignments and affect the accuracy of predictions. Second, not every biomarker is equally represented in the training data --this data imbalance is one reason why our performance on certain biomarkers is poor. We attempted to mitigate this by oversampling underrepresented biomarkers but did not observe significant improvement over equal sampling. As a result, we focus our phenotyping analysis on using the top 24 biomarkers, where performance is the most robust. Additionally, all data collected and imaged in our study was performed in-house on the same experimental setup (e.g., H&E scanner, PhenoCycler Fusion). Due to this uniformity, we have limited experimental evidence demonstrating our model’s robustness on data sourced from significantly different environments. We look forward to additional validation of our method as more paired H&E/mIF data is made publicly available.

Our study demonstrates a method for extracting multi-plex spatially resolved protein expression from H&E stains. Given the ubiquity of H&E staining in clinical workflows, a framework for enabling *in silico* staining of dozens of spatially resolved protein biomarkers offers enormous potential for improving clinical workflows and decision-making by extending the capabilities of standard histopathology.

## Materials and Methods

### CODEX data collection

All samples are prepared, stained, and acquired following CODEX User Manual Rev C (https://www.akoyabio.com).

#### Coverslip preparation

Coverslips are coated with 0.1% poly-L-lysine solution to enhance adherence of tissue sections prior to mounting. The prepared coverslips are washed and stored according to the guidelines in the CODEX User Manual.

#### Tissue sectioning

formaldehyde-fixed paraffin-embedded (FFPE) samples are sectioned at a thickness of 3-5 μm on the poly-L-lysine coated glass coverslips.

#### Antibody conjugation

Custom conjugated antibodies are prepared using the CODEX Conjugation Kit, which includes the following steps: (1) the antibody is partially reduced to expose thiol ends of the antibody heavy chains; (2) the reduced antibody is conjugated with a CODEX barcode; (3) the conjugated antibody is purified; (4) Antibody Storage Solution is added for antibody stabilization for long term storage. Post-conjugated antibodies are validated by SDS-polyacrylamide gel electrophoresis (SDS-PAGE) and quality control (QC) tissue testing, where immunofluorescence images are stained and acquired following standard CODEX protocols, then evaluated by immunologists.

#### Staining

CODEX multiplexed immunofluorescence imaging was performed on FFPE patient biopsies using the Akoya Biosciences PhenoCycler platform (also known as CODEX). 5 μm thick sections were mounted onto poly-L-lysine-treated glass coverslips as tumor microarrays. Samples were pre-treated by heating on a 55 °C hot plate for 25 minutes and cooled for 5 minutes. Each coverslip was hydrated using an ethanol series: two washes in HistoChoice Clearing Agent, two in 100% ethanol, one wash each in 90%, 70%, 50%, and 30% ethanol solutions, and two washes in deionized water (ddH2O). Next, antigen retrieval was performed by immersing coverslips in Tris-EDTA pH 9.0 and incubating them in a pressure cooker for 20 minutes on the High setting, followed by 7 minutes to cool. Coverslips were washed twice for two minutes each in ddH2O, then washed in Hydration Buffer (Akoya Biosciences) twice for two minutes each. Next, coverslips were equilibrated in Staining Buffer (Akoya Biosciences) for 30 minutes. The conjugated antibody cocktail solution in Staining Buffer was added to coverslips in a humidity chamber and incubated for 3 hours at room temperature or 16 hours at 4 °C. After incubation, the sample coverslips were washed and fixed following the CODEX User Manual.

#### Data acquisition

Sample coverslips are mounted on a microscope stage. Images are acquired using a Keyence microscope that is configured to the PhenoCycler Instrument at a 20X objective. All of the sample collections were approved by institutional review boards.

To correct for possible autofluorescence, “blank” images were acquired in each microscope channel during the first cycle of CODEX and during the last. For these images, no fluorophores were added to the tissue. These images were used for background subtraction. Typically, autofluorescence will decrease over the course of a CODEX experiment (due to repeated exposures). Thus, to correct each cycle, our method determines the extent of subtraction needed by interpolating between the first and last “blank” images.

### Sample Preprocessing

Samples are first stained and imaged using CODEX antibodies on the Akoya Biosciences PhenoCycler platform and then stained and imaged with H&E on a MoticEasyScan Pro 6N scanner with default settings and magnification. The CODEX DAPI channel and a grayscale version of the H&E image are used to perform image registration. Both images have their contrast enhanced using contrast-limited adaptive histogram equalization. The SIFT features (*35*) of each image are then found and matched based on the RANSAC algorithm (*36*) to find an image transformation from the H&E image to the CODEX coordinate space. The transformation was limited to a partial affine transformation (combinations of translation, rotation, and uniform scaling). If the initial alignment based on the grayscale H&E image was not successful, the process was repeated using the nuclear channel of the deconvolved H&E image (using a predetermined optical density matrix (*37*)), or individual color channels of the H&E image.

### Training details

The training process is visualized in Figure 1B. Each sample was first split into patches. In our standard approach, we subdivided a sample into non-overlapping 8x8px patches. The average expression within each patch is computed for every CODEX biomarker. Then, a 128 by 128px H&E patch is extracted centered on the 8x8px patch, which is used as input to the model. Thus, for instance, a 1024px by 1024px sample would yield a prediction of 128px by 128px resolution, as each 8px by 8px patch is represented by a single pixel prediction. Our model is based on a ConvNext-Small architecture with 50M parameters. The model is pre-trained on the ImageNet image dataset. During training, random augmentations are performed: horizontal and vertical flipping, brightness, contrast, saturation, hue jittering, and normalization are all performed. The training task consists of the model predicting the mean expression of the center 8x8px of a 128x128px patch for each biomarker. Thus, the model performs multitask regression (i.e., a 50-length vector) for a given patch. Model validation during training is performed using two metrics: Pearson R and SSIM (Structural Similarity Index Measure). Pearson R is a correlation metric used to assess the similarity between the measured and predicted expressions; SSIM is a similarity metric that assesses the qualitative similarity between two samples. Pearson R is computed across patches, while SSIM is computed on the reconstructed samples. This is to evaluate the model’s predictions both in terms of biological accuracy and visual similarity.

Each study has a different biomarker panel, so to account for missing biomarkers, we used a masked mean squared error loss where only the loss over present biomarkers is computed. Training is performed on 4 V100 GPUs with a batch size of 256 and a learning rate of 1e-4 with the Adam (*38*) optimizer on a schedule that reduces by half every 30K iterations. Models are trained until no improvement in this metric is observed for 75K steps.

### Sample generation

Inference is performed on the foreground H&E patches and then stitched into the predicted sample. In the standard analyses presented, a stride of 8px is used, which produces a predicted image that is 8x downsampled from the original image size. This produces predictions at a resolution of 3.02 microns per pixel). Using this image, we then produce cell-level expression predictions by upsampling the predicted image to the native resolution and then computing the average expression per cell based on the cell segmentation mask.

To produce higher-resolution images, we also demonstrate the predictions using 1px strides. In this setting, overlapping patches are generated at 64x the number of total predictions. This setting produces predictions at a resolution of 0.3775 microns per pixel. The resulting images are demonstrated and compared to the standard 9px setting in Figure S5.

### Quality control

We introduce two quantitative metrics for determining whether an H&E sample is in distribution and a high-quality generation. First, we measure the deviation between the image intensity distributions of a test H&E sample and the H&E samples in the training data. For a given test and training image pair, we extract 256 histogram bins from the image to obtain discrete distributions and then compute the Wasserstein (or Earth Mover’s) distance (also called W1 distance) between the two distributions. The quality metric for a given test image, then, is computed as the average W1 distance across all training images. By using this metric, we can *a priori* determine whether a test sample is in distribution and appropriate for evaluation.

Additionally, at the biomarker channel level, we use the dynamic range as a simple proxy for estimating the quality of a predicted sample. Since channels with very low maximum expression correlate with poor quality acquisitions (due to artifacts, staining issues, etc.), we set a threshold below which we exclude biomarkers from evaluation. Dynamic range is computed as the difference between the 99th and 1st percentile values in a generated biomarker stain. Since the dynamic range is only a function of the predicted image, it does not depend on having ground truth CODEX measurements for an H&E sample.

### Expression Metrics

We report three primary evaluation metrics for patch-level predictions: Pearson R (<u>sklearn.metrics.pearsonr</u>), Spearman R (<u>sklearn.metrics.spearmanr</u>), and concordance index, or C-index (<u>lifelines.utils.concordance_index</u>). Pearson correlation is a measure of the linear relationship between the ground truth and predicted biomarker expressions. Additionally, we report two rank metrics (Spearman R and C-index) to assess the model predictions’ usefulness in clinical tasks that rely on ordering patient samples by a specific biomarker expression or cell type count. Pearson and Spearman correlations are calculated as the average correlations across all ground truth and predicted CODEX patches. C-index is computed on the 75th percentile values for both ground truth and predicted CODEX, for each biomarker and across all samples. Only during training, SSIM is computed across ground truth and predicted images. To ensure that the metrics are calculated on valid data points, we exclude patches with a lower groundtruth expression value than the 90th percentile value of background noise for each biomarker.

### Baseline Methods

We also introduce several baseline methods for comparison to *ROSIE*:

First, we use the H&E sample alone to predict CODEX expression (called *H&E expression)*. In this method, we apply a simple threshold (>50), averaged across the three color channels, and then use the intensity to predict each biomarker. This is to evaluate the similarity of the hematoxylin and eosin stains to the CODEX stains and to validate that the model is not simply recapitulating stain intensity in its CODEX predictions.

Second, we compute morphology statistics based on the segmentation masks calculated from the DAPI channel (called *cell morphology)*. Cell segmentation is performed using the DeepCell algorithm (*39*). We use the HistomicsTK <u>compute_morphometry_features</u> function and extract 19 features in total: Orientation, Area, Convex Hull Area, Major Axis Length, Minor Axis Length, Perimeter, Circularity, Eccentricity, Equivalent Diameter, Extent, Minor to Major Axis Ratio, Solidity, Hu Moments (1st to 7th). In addition, we compute the average across each of the RGB channels and include these as three additional features. We train a three-layer multi-layer perceptron neural network using these features as input to predict the expression of 50 protein biomarkers. Each layer has 100 nodes followed by a ReLU activation function and is trained with a 1e-4 learning rate, mean squared error loss, and Adam optimizer. The weights that generated the best validation accuracy after 50 epochs are used. This is a stronger baseline that is intended to represent typical features (morphology and intensity) derived from H&E images.

Finally, as an additional baseline for cell phenotyping, we assign cell labels randomly according to the average ground truth cell label proportions across all samples (called *bulk phenotyping).* This method approximates estimating cell type proportions through a cheaper, more readily available phenotyping technique than CODEX (like flow cytometry) and then using them to infer spatially located cell types.

### Cell Phenotyping

Cell phenotyping metrics are computed using the following steps: First, cell clusters are produced using Leiden clustering based on the cell-level CODEX measured expressions. These clusters are identified and merged based on cell expression within each cluster to produce manually annotated cell labels. Then, we trained a k-nearest neighbors algorithm (where k=100) to generate a graph based on these clusters, which is used to automatically generate the reference cell labels. We use the same kNN and the predicted expressions to generate the predicted cell labels. We report the F1 scores, which are relative to the cell typing determined using clustering on the *CODEX* measurements. In this analysis, we use only the top 24 biomarkers by Pearson correlation as input features to the kNN algorithm. Figure S7 shows the mean biomarker expressions for each defined cell type. For cell phenotyping performed on Ochsner-CRC, we similarly train a kNN on manually annotated cell labels and use it to generate reference cell labels. For uniformity of comparision, we define the same cell phenotypes as in the Stanford-PGC dataset.

### Tissue Structure Discovery

The Spatial Cellular Graph Partitioning (SCGP) framework is described in full detail in (*33*). The algorithm is summarized in the following steps:

1. Construct a graph with cells as nodes. Spatial edges are added between neighboring cell pairs, and feature edges are added for cell pairs with similar expression profiles.
2. Partitions are detected by community detection algorithms such as the Leiden algorithm (*40*).
3. Each partition is manually annotated based on its underlying expression profile and cell morphology.

The above steps are performed independently on the ground truth and H&E imputed mIF samples, and a mapping from the imputed partitions to the ground truth partitions is calculated. Finally, we compute the adjusted Rand Index (ARI) and F1 scores. ARI measures the similarity between the ground truth and imputation-derived partitions and does not require cluster labeling; the F1 score is computed over manually annotated labels. As a baseline, we perform the partitioning over morphology features extracted from the cell segmentations, as well as adding in the average RGB expressions per cell. Figure S7 shows the mean biomarker expressions for each defined tissue structure.

### Cell neighborhood phenotyping

To define cell neighbors, we first perform cell segmentation on the DAPI channel for each sample. Based on the computed cell centroids, we then construct a Delauney triangulation and Voronoi diagram, from which we then construct a graph with cells as nodes and Delauney neighbors as edges. To define lymphocyte neighboring epithelial cells, we identify epithelial cells and then find the subset of these cells that share an edge with a lymphocyte (B cell or T cell). The reported percentage of LNEs is defined as the proportion of epithelial cells in a sample that are LNEs. Additionally, we define tumor-infiltrating lymphocytes (TILs) as lymphocytes embedded in tumor regions. To identify TILs, we find lymphocytes that are assigned to epithelial tissue structures. For each sample, we report the raw count of TILs.

## Author Contributions

Conceptualization: EW, AET, ATM, JZ

Methodology, Investigation, and Visualization: EW, AET, ZW, NT

Resources: GWC, AM, CS, JRH, UP, CIL, LL, HG, VB, AR

Supervision: AET, ATM, JZ Writing—original draft: EW, AET

Writing—review & editing: MB, ZW, ATM, JZ

## Data and Materials Availability

The code and data used to produce the analysis for this manuscript are available at https://gitlab.com/enable-medicine-public/rosie. All data are available in the main text or the supplementary materials.

## Supplementary Materials

**Supplemental Table 1:** Comparison of model architectures. Our CNN-based approach outperformed larger Vision Transformer models and histopathology image pre-training approaches.

| Stanford-PGC<br>(50 biomarkers) | Model<br>parameters (#) | Pearson R | Spearman R | C-index (sample) |
| --- | --- | --- | --- | --- |
| - *ConvNext*<br>(Ours) | 50.2M | 0.319 | 0.386 | 0.694 |
| - UNI<br>(VitL-16) | 304.3M | 0.278 | 0.337 | 0.619 |
| - VitB-16 | 86.6M | 0.294 | 0.356 | 0.626 |

**Supplementary Figure 1:**
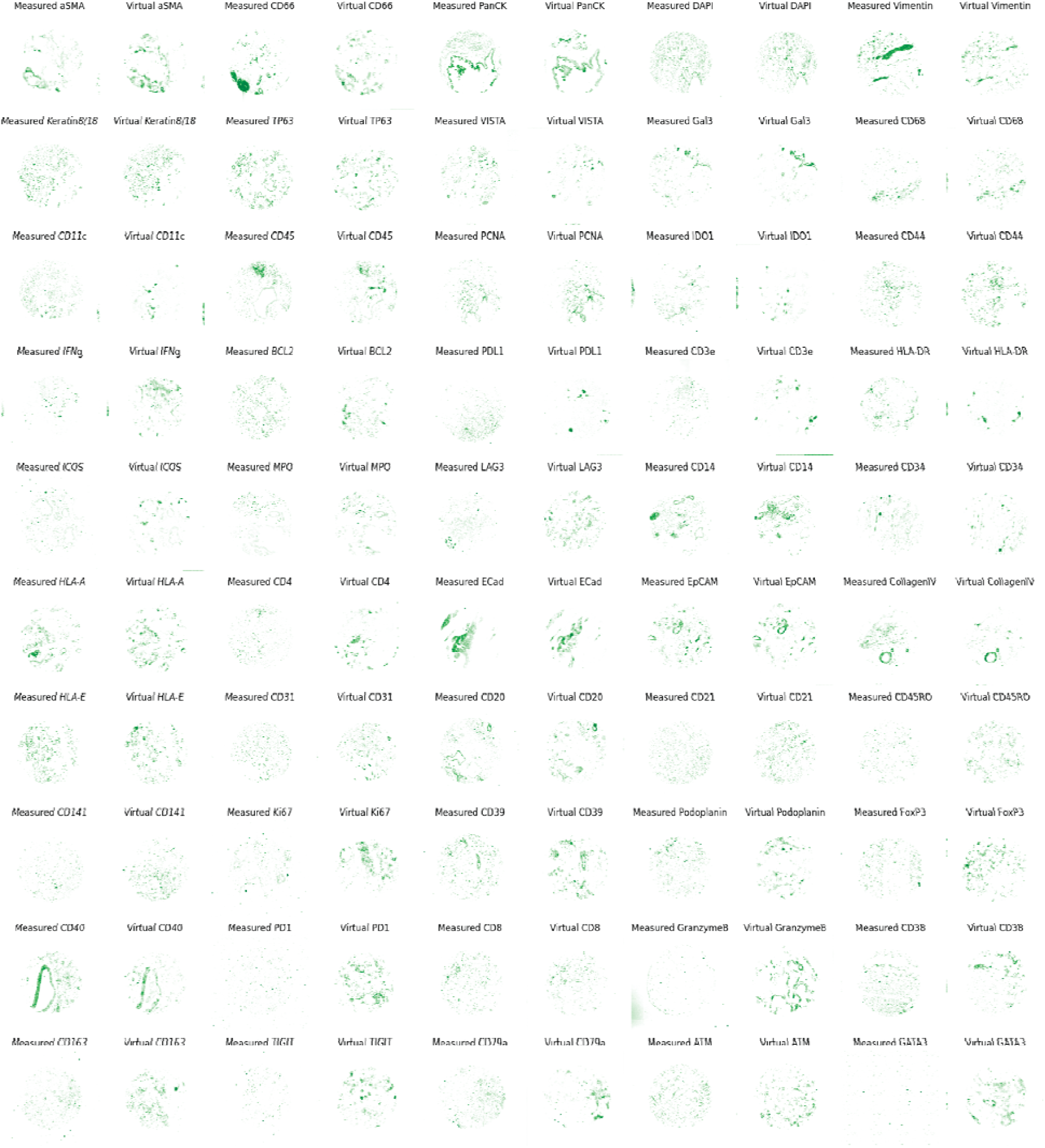
Median samples (by Pearson R) are visualized for all 50 biomarkers from the Stanford-PGC test set.

**Supplementary Figure 2:**
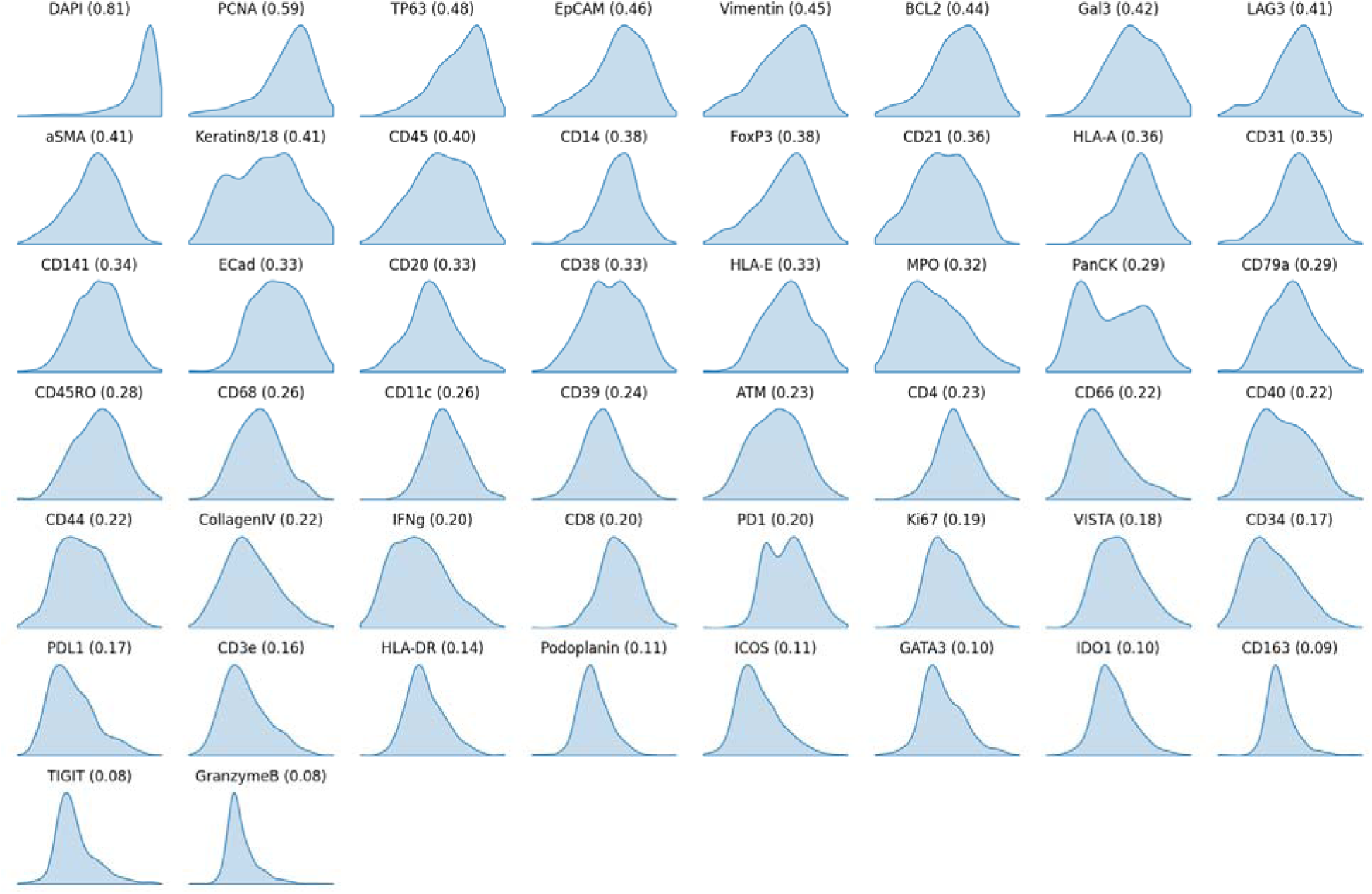
Density plot of Pearson R correlation for each biomarker across the distribution of samples in the Stanford-PGC test set.

**Supplementary Figure 3:**
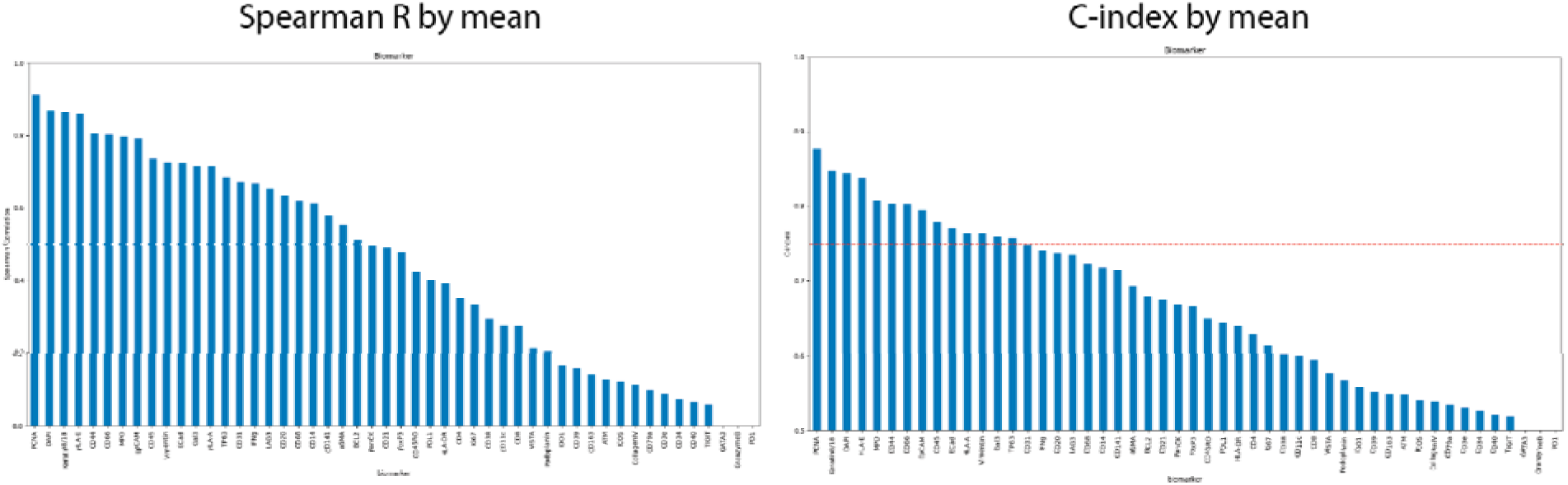
Rank tests on each biomarker comparing ground truth and *ROSIE*-generated expressions in the Stanford-PGC test set.

**Supplementary Table 2:**
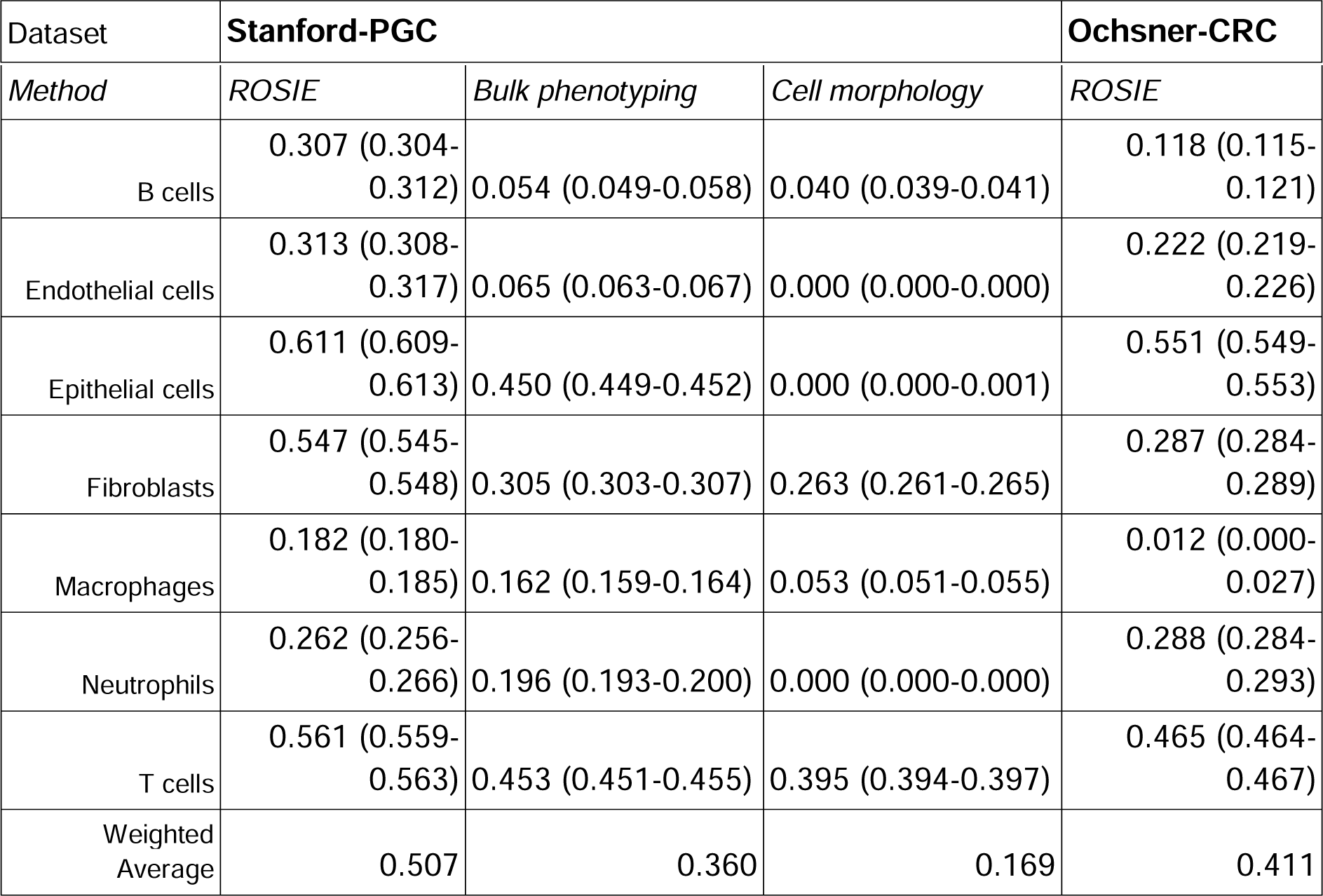
Cell type predictions by individual cell labels on the Stanford-PGC and Ochsner-CRC test datasets, along with two baseline methods (Bulk phenotyping and Cell morphology). We report F1 scores for each cell label and 95% bootstrapped confidence intervals in parentheses.

**Supplementary Figure 4:**
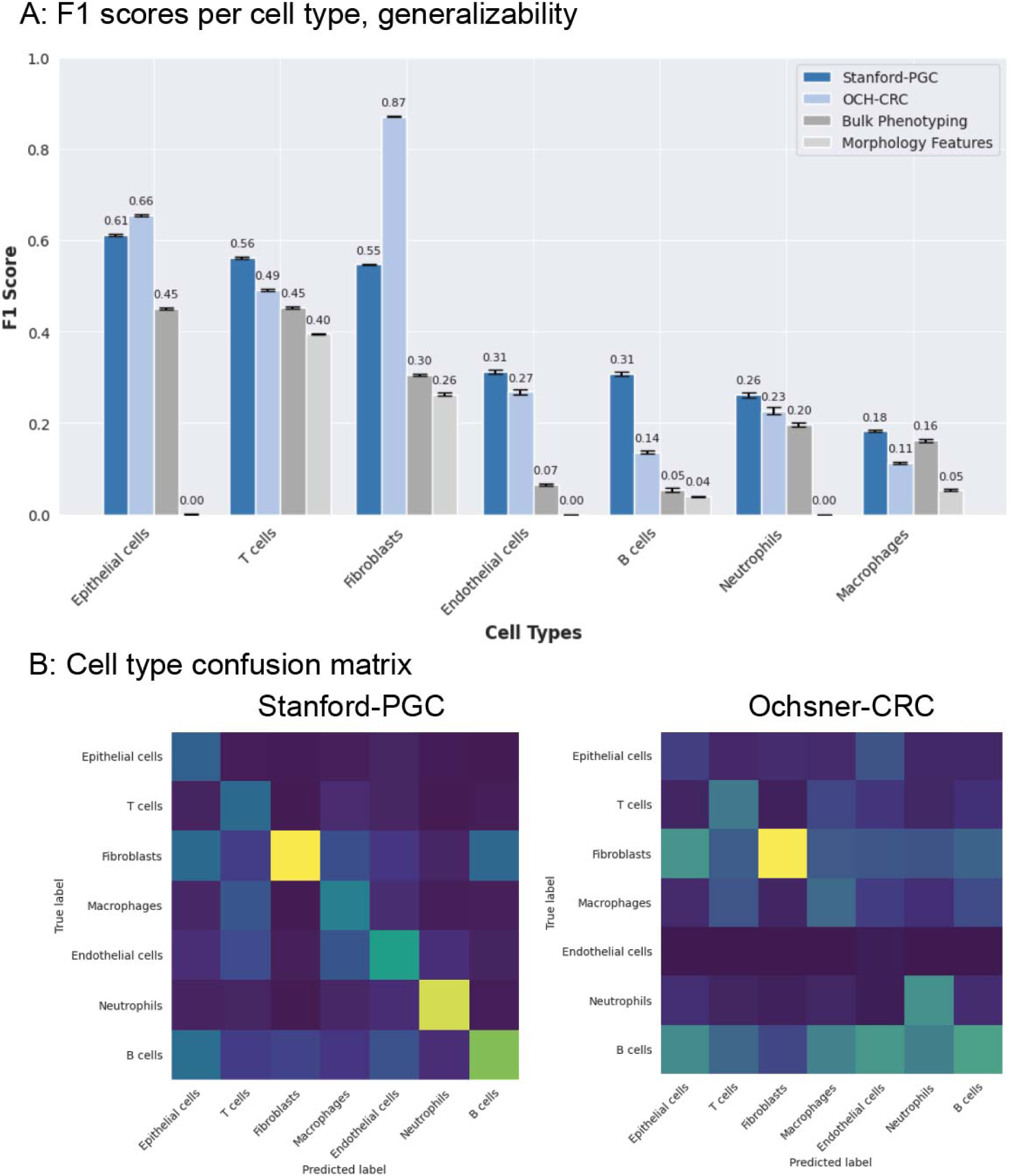
*ROSIE* generalizes well to predicting cell types in a study of not-seen-before colorectal cancer tumor samples (Ochsner-CRC). **A:** The cell type predictions on Ochsner-CRC are comparable to those in Stanford-PGC and two baseline methods, Bulk Phenotyping and Morphology Features. **B:** Cell type confusion matrices for predictions from *ROSIE* on the Stanford-PGC and Ochsner-CRC datasets. Of note is that B cells and T cells are well differentiated in cell predictions.

**Supplemental Figure 5:**
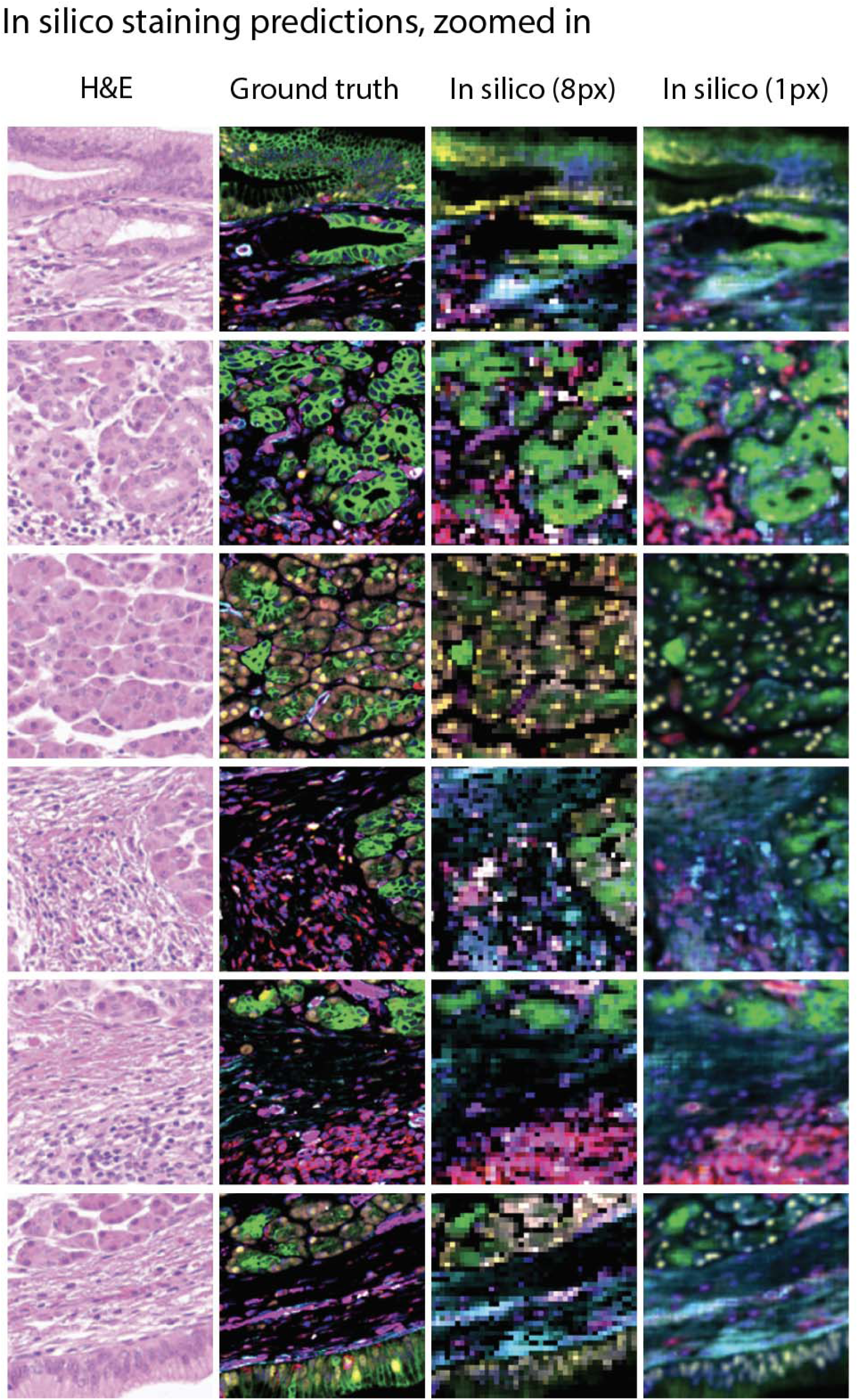
Zoomed in patches of predictions from *ROSIE*, using an 8px striding window and 1px striding window. With 1px, the resolution and detail are significantly improved; however, generating a sample requires 64x the amount of compute resources. All analyses in this study are performed on the 8px sliding window-generated images.

**Supplemental Figure 6:**
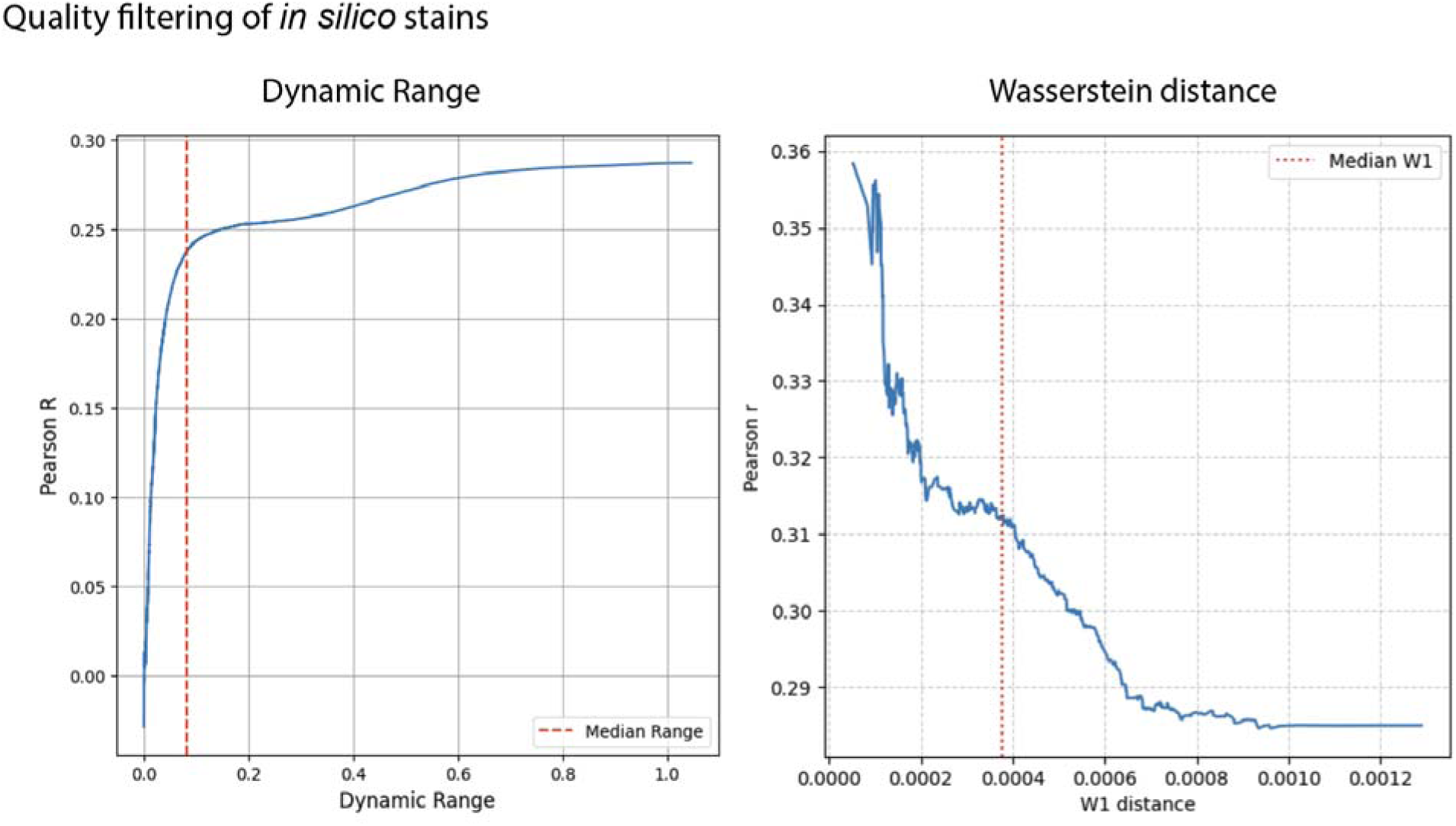
Quality filtering: We show two simple methods for predicting the quality of a predicted stain using *ROSIE*. **Left**: Low-quality predictions yield samples with low dynamic range; based on this intuition, we show that the predictive accuracy (measured by average Pearson R) of a biomarker stain is strongly positively correlated with a higher dynamic range. **Right**: We also compute the Wasserstein distance between the H&E image intensity histograms of the training and test data. We also find that higher W1 distances are associated with lower Pearson R scores. In both plots, each point represents the average Pearson R of all samples with measured values less than or equal to the value on the x-axis.

**Supplementary Figure 7:**
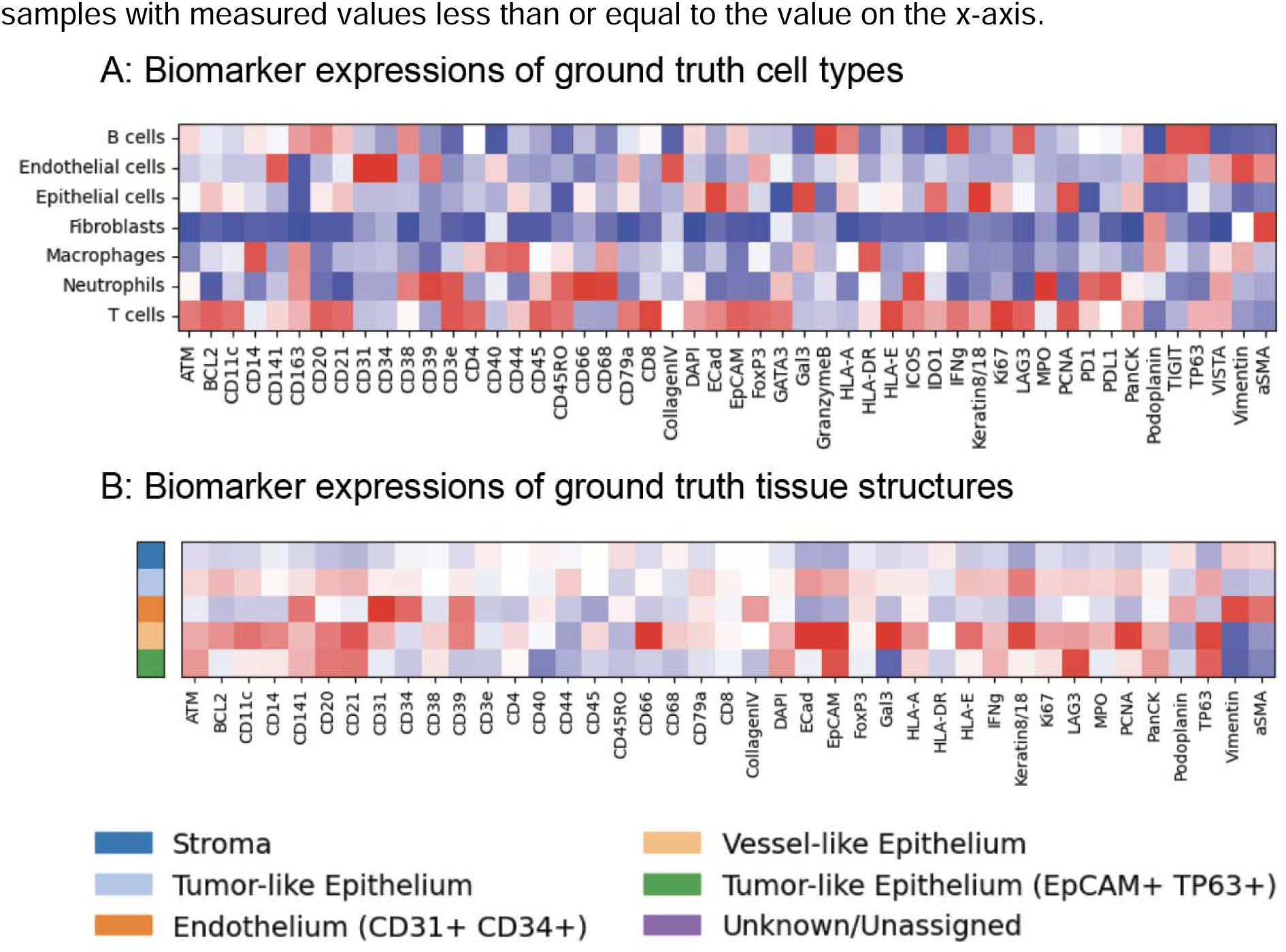
A: The mean expression profile for each ground truth cell phenotype on the Stanford-PGC test dataset. Each biomarker expression is mean normalized across the cell types. **B:** The mean expression profile for each ground truth tissue structure on the Stanford-PGC test dataset.

**Supplementary Figure 8:**
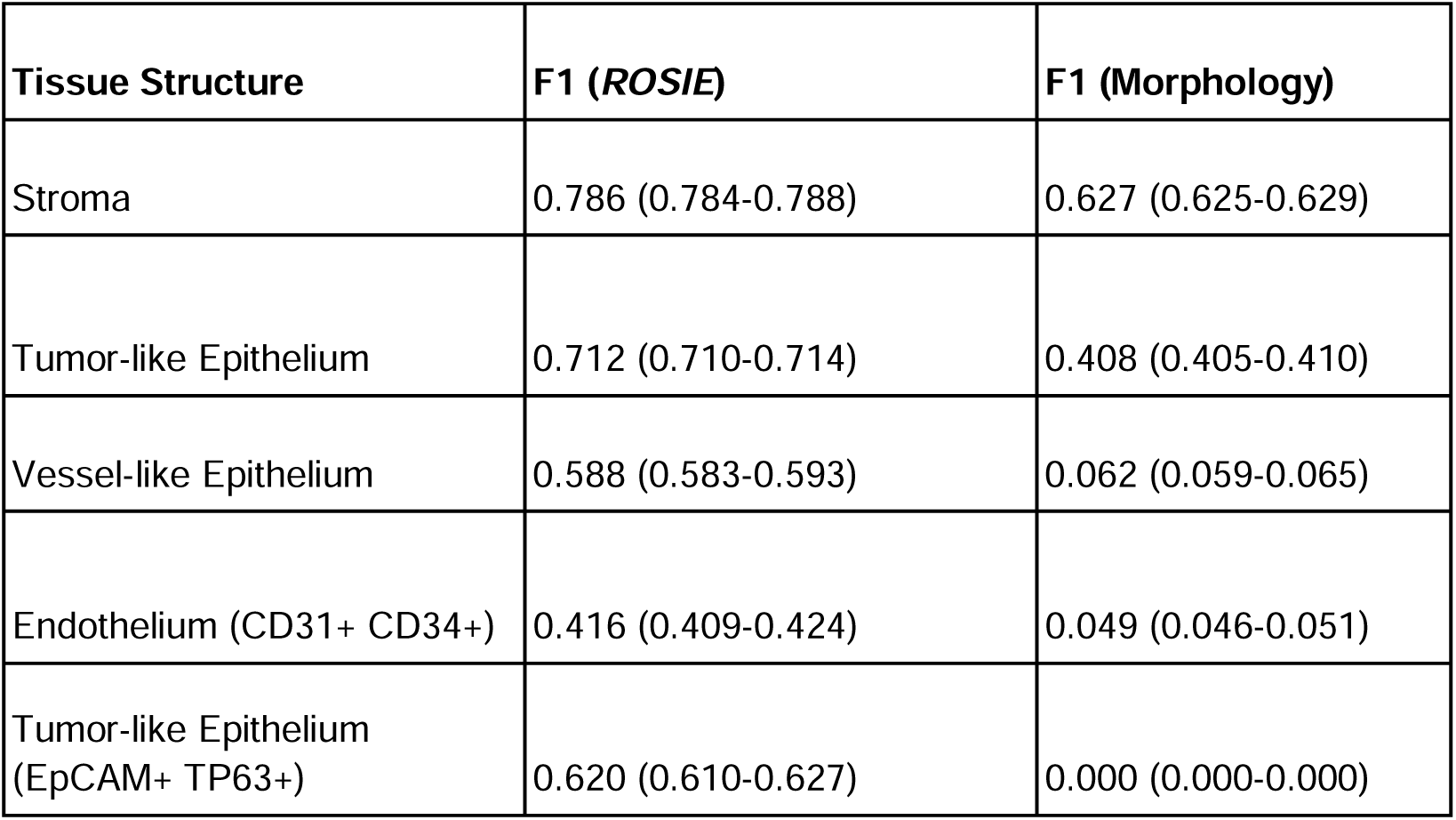
Tissue structure discovery performance by individual tissue structure type. 95% bootstrapped confidence intervals are reported in parentheses.

## References

1. H. D. Couture, Deep learning-based prediction of molecular tumor biomarkers from H&E: A practical review. J. Pers. Med. 12, 2022 (2022).

2. S. Graham, Q. D. Vu, S. E. A. Raza, A. Azam, Y. W. Tsang, J. T. Kwak, N. Rajpoot, HoVer-net: Simultaneous segmentation and classification of nuclei in multi-tissue histology images, arXiv [cs.CV] (2018). http://arxiv.org/abs/1812.06499.

3. J. Gamper, N. Alemi Koohbanani, K. Benet, A. Khuram, N. Rajpoot, “PanNuke: An Open Pan-Cancer Histology Dataset for Nuclei Instance Segmentation and Classification” in Digital Pathology (Springer International Publishing, Cham, 2019)*Lecture notes in computer science*, pp. 11–19.

4. M. Amgad, L. A. Atteya, H. Hussein, K. Mohammed, E. Hafiz, M. A. T. Elsebaie, Alhusseiny, M. A. AlMoslemany, A. M. Elmatboly, P. A. Pappalardo, R. Sakr, P. Mobadersany, A. Rachid, A. M. Saad, A. Alkashash, I. A. Ruhban, A. Alrefai, N. M. Elgazar, A. Abdulkarim, A.-A. Farag, A. Etman, A. G. Elsaeed, Y. Alagha, Y. A. Amer, A. Raslan, M. K. Nadim, M. A. T. Elsebaie, A. A. Ayad, L. E. Hanna, A. Gadallah, M. Elkady, Drumheller, D. Jaye, D. Manthey, D. Gutman, H. Elfandy, Lee A. D. Cooper Department of Pathology, Northwestern University, Chicago., Il, Usa, Cairo Health Care Administration, Egyptian Ministry of Health, Cairo, Egypt., D. Pathology, Nasser institute for research, Treatment, Laboratory Medicine, U. Pennsylvania, Pa, Department of Clinical Research, the Bartol Research Institute, Giza, Department of Preventive Medicine, Cook County Hospital, Baystate Medical Center, U. Massachusetts, Springfield, Ma., Faculty of Veterinary Medicine, M. University, Menoufia, Al-Azhar University, Consultant for The Center for Applied Proteomics, Molecular Medicine, George Mason University, Manassas, Va, National Liver Institute, Ain Shams University, Cleveland Clinic Foundation, Cleveland, Oh, I. University, Indianapolis, In, Damascus University, Damascus, Syria, M. University, Mansoura, Cairo University, Department of Anaesthesia, Critical Care, Menoufia University Hospital, D. Pathology, R. Department, Oncology Consultants, Houston, Tx, S. D. Informatics, Pine Brook, Nj, Emory University School of Medicine, Atlanta, Ga, Kitware Inc., C. Park, Ny, D. Neurology, National Cancer Institute, Children’s Cancer Hospital Egypt Cche, Lurie Cancer Center, Center for Computational Imaging, S. Analytics, Northwestern University Feinberg School of Medicine, NuCLS: A scalable crowdsourcing approach and dataset for nucleus classification and segmentation in breast cancer. Gigascience 11 (2021).

5. J. W. Hickey, Y. Tan, G. P. Nolan, Y. Goltsev, Strategies for accurate cell type identification in CODEX multiplexed imaging data. Front. Immunol. 12, 727626 (2021).

6. Y. Goltsev, N. Samusik, J. Kennedy-Darling, S. Bhate, M. Hale, G. Vazquez, S. Black, G. P. Nolan, Deep Profiling of Mouse Splenic Architecture with CODEX Multiplexed Imaging. Cell 174, 968–981.e15 (2018).

7. Z. Wu, A. E. Trevino, E. Wu, K. Swanson, H. J. Kim, H. B. D’Angio, R. Preska, G. W. Charville, P. D. Dalerba, A. M. Egloff, R. Uppaluri, U. Duvvuri, A. T. Mayer, J. Zou, Graph deep learning for the characterization of tumour microenvironments from spatial protein profiles in tissue specimens. Nat Biomed Eng 6, 1435–1448 (2022).

8. D. Phillips, C. M. Schürch, M. S. Khodadoust, Y. H. Kim, G. P. Nolan, S. Jiang, Highly multiplexed phenotyping of immunoregulatory proteins in the tumor microenvironment by CODEX tissue imaging. Front. Immunol. 12, 687673 (2021).

9. S. Black, D. Phillips, J. W. Hickey, J. Kennedy-Darling, V. G. Venkataraaman, N. Samusik, Y. Goltsev, C. M. Schürch, G. P. Nolan, CODEX multiplexed tissue imaging with DNA-conjugated antibodies. Nat. Protoc. 16, 3802–3835 (2021).

10. Z. Huang, F. Bianchi, M. Yuksekgonul, T. J. Montine, J. Zou, A visual-language foundation model for pathology image analysis using medical Twitter. Nat. Med. 29, 2307– 2316 (2023).

11. M. Y. Lu, B. Chen, D. F. K. Williamson, R. J. Chen, I. Liang, T. Ding, G. Jaume, I. Odintsov, L. P. Le, G. Gerber, A. V. Parwani, A. Zhang, F. Mahmood, A visual-language foundation model for computational pathology. Nat. Med. 30, 863–874 (2024).

12. R. J. Chen, T. Ding, M. Y. Lu, D. F. K. Williamson, G. Jaume, A. H. Song, B. Chen, A. Zhang, D. Shao, M. Shaban, M. Williams, L. Oldenburg, L. L. Weishaupt, J. J. Wang, A. Vaidya, L. P. Le, G. Gerber, S. Sahai, W. Williams, F. Mahmood, Towards a general-purpose foundation model for computational pathology. Nat. Med. 30, 850–862 (2024).

13. P. Pati, S. Karkampouna, F. Bonollo, E. Compérat, M. Radic, M. Spahn, A. Martinelli, M. Wartenberg, M. K. Julio, M. A. Rapsomaniki, Multiplexed tumor profiling with generative AI accelerates histopathology workflows and improves clinical predictions, bioRxiv (2023)p. 2023.11.29.568996.

14. S. Andani, B. Chen, J. Ficek-Pascual, S. Heinke, R. Casanova, B. Sobottka, B. Bodenmiller, Tumor Profiler Consortium Viktor H. Koelzer, G. Rätsch, Multi-V-Stain: Multiplexed Virtual Staining of Histopathology Whole-Slide Images. doi: 10.1101/2024.01.26.24301803.

15. P. Ghahremani, Y. Li, A. Kaufman, R. Vanguri, N. Greenwald, M. Angelo, T. J. Hollmann, S. Nadeem, Deep Learning-Inferred Multiplex ImmunoFluorescence for Immunohistochemical Image Quantification. Nat Mach Intell 4, 401–412 (2022).

16. E. A. Burlingame, M. McDonnell, G. F. Schau, G. Thibault, C. Lanciault, T. Morgan, B. E. Johnson, C. Corless, J. W. Gray, Y. H. Chang, SHIFT: speedy histological-to-immunofluorescent translation of a tumor signature enabled by deep learning. Sci. Rep. 10, 17507 (2020).

17. N. Bouteldja, D. L. Hölscher, R. D. Bülow, I. S. D. Roberts, R. Coppo, P. Boor, Tackling stain variability using CycleGAN-based stain augmentation. J. Pathol. Inform. 13, 100140 (2022).

18. N. Bouteldja, B. M. Klinkhammer, T. Schlaich, P. Boor, D. Merhof, Improving unsupervised stain-to-stain translation using self-supervision and meta-learning. J. Pathol. Inform. 13, 100107 (2022).

19. H. Wieslander, A. Gupta, E. Bergman, E. Hallström, P. J. Harrison, Learning to see colours: generating biologically relevant fluorescent labels from bright-field images, bioRxiv (2021)p. 2021.01.18.427121.

20. O. Cetin, M. Chen, P. Ziegler, P. Wild, H. Koeppl, “Deep learning-based restaining of histopathological images” in 2022 IEEE International Conference on Bioinformatics and Biomedicine (BIBM) (IEEE, 2022), pp. 1467–1474.

21. K. de Haan, Y. Zhang, J. E. Zuckerman, T. Liu, A. E. Sisk, M. F. P. Diaz, K.-Y. Jen, A. Nobori, S. Liou, S. Zhang, R. Riahi, Y. Rivenson, W. D. Wallace, A. Ozcan, Deep learning-based transformation of H&E stained tissues into special stains. Nat. Commun. 12, 4884 (2021).

22. B. He, S. Bukhari, E. Fox, A. Abid, J. Shen, C. Kawas, M. Corrada, T. Montine, J. Zou, AI-enabled in silico immunohistochemical characterization for Alzheimer’s disease. Cell Rep Methods 2, 100191 (2022).

23. C. Bian, B. Philips, T. Cootes, M. Fergie, HEMIT: H&E to Multiplex-immunohistochemistry Image Translation with Dual-Branch Pix2pix Generator, arXiv [eess.IV] (2024). http://arxiv.org/abs/2403.18501.

24. G. Srinivasan, M. Davis, M. LeBoeuf, M. Fatemi, Z. Azher, Y. Lu, A. Diallo, M. S. Montivero, F. Kolling, L. Perrard, L. Salas, B. Christensen, S. Palisoul, G. Tsongalis, L. Vaickus, S. Preum, J. Levy, Potential to Enhance Large Scale Molecular Assessments of Skin Photoaging through Virtual Inference of Spatial Transcriptomics from Routine Staining. bioRxiv, doi: 10.1101/2023.07.30.551188 (2023).

25. E. Wu, A. E. Trevino, Z. Wu, K. Swanson, H. J. Kim, H. B. D’Angio, R. Preska, A. E. Chiou, G. W. Charville, P. Dalerba, U. Duvvuri, A. D. Colevas, J. Levi, N. Bedi, S. Chang, J. Sunwoo, A. M. Egloff, R. Uppaluri, A. T. Mayer, J. Zou, 7-UP: Generating in silico CODEX from a small set of immunofluorescence markers. PNAS Nexus 2, gad171 (2023).

26. Z. Zhou, Y. Jiang, Z. Sun, T. Zhang, W. Feng, G. Li, R. Li, L. Xing, Virtual multiplexed immunofluorescence staining from non-antibody-stained fluorescence imaging for gastric cancer prognosis. EBioMedicine 107, 105287 (2024).

27. Z. Liu, H. Mao, C.-Y. Wu, C. Feichtenhofer, T. Darrell, S. Xie, A ConvNet for the 2020s, arXiv [cs.CV] (2022). http://arxiv.org/abs/2201.03545.

28. A. Dosovitskiy, L. Beyer, A. Kolesnikov, D. Weissenborn, X. Zhai, T. Unterthiner, M. Dehghani, M. Minderer, G. Heigold, S. Gelly, J. Uszkoreit, N. Houlsby, An image is worth 16x16 words: Transformers for image recognition at scale, arXiv [cs.CV] (2020). http://arxiv.org/abs/2010.11929.

29. F. M. Howard, J. Dolezal, S. Kochanny, J. Schulte, H. Chen, L. Heij, D. Huo, R. Nanda, O. I. Olopade, J. N. Kather, N. Cipriani, R. L. Grossman, A. T. Pearson, The impact of site-specific digital histology signatures on deep learning model accuracy and bias. Nat. Commun. 12, 4423 (2021).

30. M. Schmitt, R. C. Maron, A. Hekler, A. Stenzinger, A. Hauschild, M. Weichenthal, M. Tiemann, D. Krahl, H. Kutzner, J. S. Utikal, S. Haferkamp, J. N. Kather, F. Klauschen, E. Krieghoff-Henning, S. Fröhling, C. von Kalle, T. J. Brinker, Hidden variables in deep learning digital pathology and their potential to cause batch effects: Prediction model study. J. Med. Internet Res. 23, e23436 (2021).

31. P. Ouyang, L. Wang, J. Wu, Y. Tian, C. Chen, D. Li, Z. Yao, R. Chen, G. Xiang, J. Gong, Z. Bao, Overcoming cold tumors: a combination strategy of immune checkpoint inhibitors. Front. Immunol. 15, 1344272 (2024).

32. C. Hartupee, B. M. Nagalo, C. Y. Chabu, M. Z. Tesfay, J. Coleman-Barnett, J. T. West, O. Moaven, Pancreatic cancer tumor microenvironment is a major therapeutic barrier and target. Front. Immunol. 15, 1287459 (2024).

33. Z. Wu, A. Kondo, M. McGrady, E. A. G. Baker, B. Chidester, E. Wu, M. K. Rahim, N. A. Bracey, V. Charu, R. J. Cho, J. B. Cheng, M. Afkarian, J. Zou, A. T. Mayer, A. E. Trevino, Discovery and generalization of tissue structures from spatial omics data. Cell Rep. Methods 4, 100838 (2024).

34. X. Wang, J. Zhao, E. Marostica, W. Yuan, J. Jin, J. Zhang, R. Li, H. Tang, K. Wang, Y. Li, F. Wang, Y. Peng, J. Zhu, J. Zhang, C. R. Jackson, J. Zhang, D. Dillon, N. U. Lin, L. Sholl, T. Denize, D. Meredith, K. L. Ligon, S. Signoretti, S. Ogino, J. A. Golden, M. P. Nasrallah, X. Han, S. Yang, K.-H. Yu, A pathology foundation model for cancer diagnosis and prognosis prediction. Nature, 1–9 (2024).

35. D. G. Lowe, Distinctive image features from scale-invariant keypoints. Int. J. Comput. Vis. 60, 91–110 (2004).

36. M. A. Fischler, R. C. Bolles, Random sample consensus: a paradigm for model fitting with applications to image analysis and automated cartography. Commun. ACM 24, 381–395 (1981).

37. A. C. Ruifrok, D. A. Johnston, Quantification of histochemical staining by color deconvolution. Anal. Quant. Cytol. Histol. 23, 291–299 (2001).

38. D. P. Kingma, J. Ba, Adam: A method for stochastic optimization, arXiv [cs.LG] (2014). http://arxiv.org/abs/1412.6980.

39. N. F. Greenwald, G. Miller, E. Moen, A. Kong, A. Kagel, T. Dougherty, C. C. Fullaway, B. J. McIntosh, K. X. Leow, M. S. Shwartz, C. Pavelchek, S. Cui, I. Camplisson, O. Bar-Tal, J. Singh, M. Fong, G. Chaudhry, Z. Abraham, J. Moseley, S. Warshawsky, E. Soon, S. Greenbaum, T. Risom, T. Hollmann, S. C. Bendall, L. Keren, W. Graf, M. Angelo, D. Van Valen, Whole-cell segmentation of tissue images with human-level performance using large-scale data annotation and deep learning. Nat. Biotechnol. 40, 555–565 (2022).

40. V. A. Traag, L. Waltman, N. J. van Eck, From Louvain to Leiden: guaranteeing well-connected communities. Sci. Rep. 9, 5233 (2019).

